# ISSAAC-seq enables sensitive and flexible multimodal profiling of chromatin accessibility and gene expression in single cells

**DOI:** 10.1101/2022.01.16.476488

**Authors:** Wei Xu, Weilong Yang, Yunlong Zhang, Yawen Chen, Ni Hong, Qian Zhang, Xuefei Wang, Yukun Hu, Kun Song, Wenfei Jin, Xi Chen

## Abstract

Joint profiling of chromatin accessibility and gene expression from the same single cell/nucleus provides critical information about cell types in a tissue and cell states during a dynamic process. These emerging multi-omics techniques help the investigation of cell-type resolved gene regulatory mechanisms^1–7^. However, many methods are currently limited by low sensitivity, low throughput or complex workflow. Here, we developed in situ SHERRY after ATAC-seq (ISSAAC-seq), a highly sensitive and flexible single cell multi-omics method to interrogate chromatin accessibility and gene expression from the same single nucleus. We demonstrated that ISSAAC-seq is sensitive and provides high quality data with orders of magnitude more features than existing methods. Using the joint profiles from over 10,000 nuclei from the mouse cerebral cortex, we uncovered major and rare cell types and cell-type specific regulatory elements and identified heterogeneity at the chromatin level within established cell types defined by gene expression. Finally, we revealed distinct dynamics and relationships of gene expression and chromatin accessibility during an oligodendrocyte maturation trajectory.

## Main Text

Recent technological development enables detailed characterization of various modalities at the single cell level, such as gene expression^8–13^, chromatin accessibility^14–20^ and protein abundance^21,22^. Data from different modalities provide distinct and complementary information about cell types or states. Currently, most methods only assay one modality of the cell at a time. Though highly informative, only one molecular layer of the cell is profiled, which is insufficient to provide comprehensive views of cell states or the associated regulatory programs^23,24^. Certain computational methods allow the integration of different molecular layers during downstream analysis. However, the performance of such integration is difficult to assess^23,25^. In addition, many integration methods are predicated on the assumption that data from different modalities can be projected to some shared latent space, which remains to be rigorously tested^23,25^. To circumvent these limitations, new methods continue to be developed for the parallel detection of different molecular layers from the same single cell^1–7,26–31^.

Here we developed ISSAAC-seq, a highly sensitive and scalable method to interrogate chromatin accessibility and gene expression from the same single nucleus with a flexible workflow. The method is based on the combination of the recently developed Sequencing HEteRo RNA-DNA-hYbrid (SHERRY)^32,33^ and previously established scATAC-seq approaches^16,19,20,34^ (**Fig. 1a, Supplementary Fig. 1 and 2**). In ISSAAC-seq, open chromatin regions are tagged by the transposase Tn5 with regular Nextera adaptors that are used in a typical scATAC-seq experiment. Then a primer containing a partial TruSeq adaptor, a 10-bp unique molecular identifier (UMI) and poly-T is added for the reverse transcription in the nucleus^7,35^. Subsequently, a Tn5 homodimer loaded with one side of the Nextera sequence is added to mark the RNA/DNA hybrid^32,33^. Up to this stage, every step is performed in the nucleus (*in situ*) of a population of cells without physical separation of chromatin and RNA. Finally, single nuclei can be isolated for cell barcode addition and library preparation in different ways, such as flow cytometry (FACS) for limited cell number and droplet-based microfluidics for high throughput studies (**Fig. 1a**). After cell pooling and pre-amplification, the library is split into two equal portions. One portion is amplified by a Nextera primer pair for chromatin accessibility sequencing (ATAC-seq), and the other portion is amplified with a Nextera and a TruSeq primer pair for gene expression sequencing (RNA-seq) (**Fig. 1a, Supplementary Fig. 1** and **2**).

**Fig. 1.**
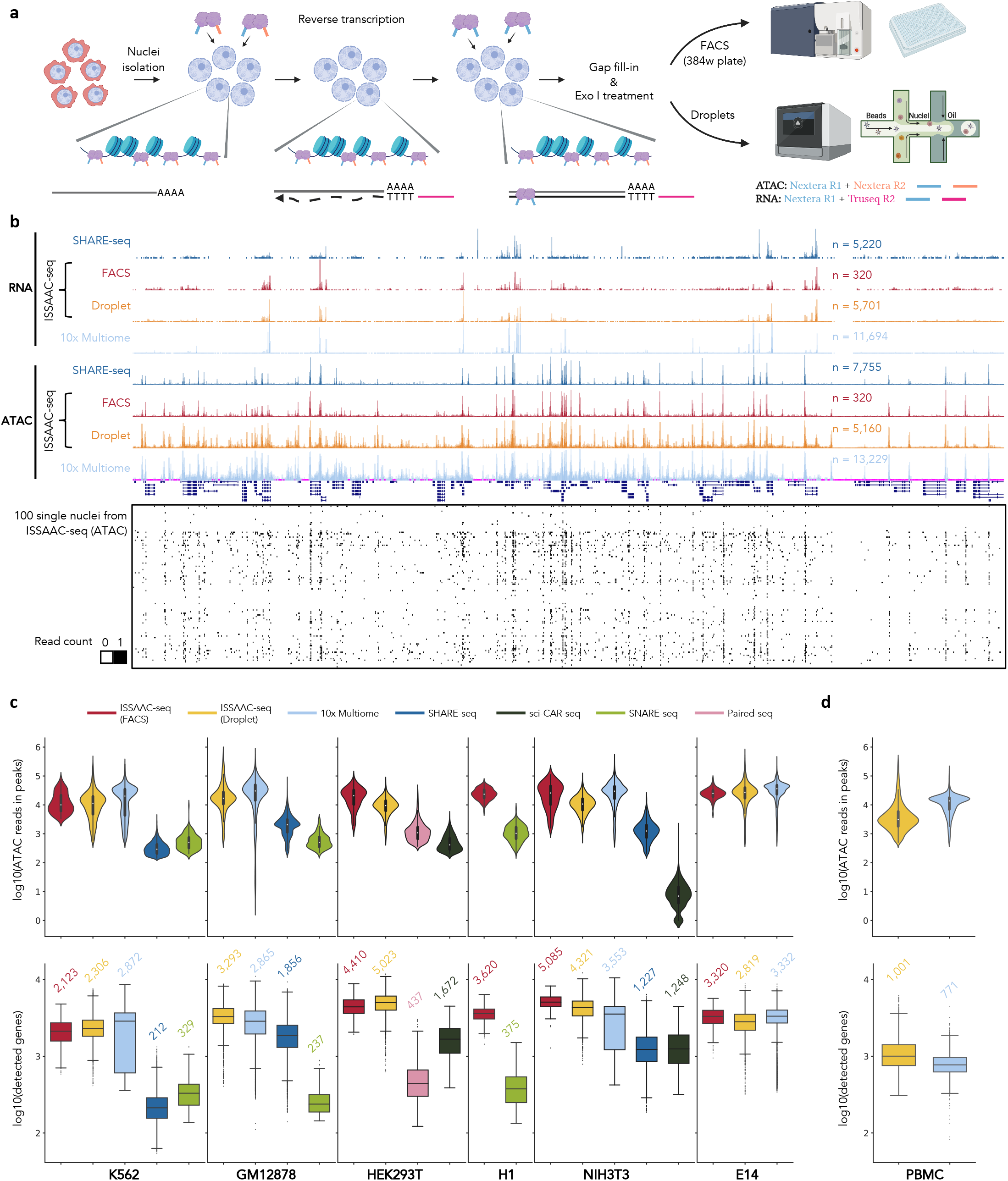
Joint profiling of chromatin accessibility and gene expression using ISSAAC-seq. **a**, A schematic view of the ISSAAC-seq workflow. Key steps and adaptor configurations are outlined in the main text and Supplementary Figs. 1 & 2. **b**, UCSC genome browser tracks showed the aggregated signal of single cells (ATAC and RNA) of K562 cells from SHARE-seq, ISSAAC-seq and the 10x Multiome kit. ATAC-seq signals from 100 randomly selected single cells were shown at the bottom. **c** and **d**, Distributions of the numbers of reads in peaks (ATAC) and detected genes (RNA) from different methods performed in the indicated cell lines (**c**) or human PBMCs (**d**). Median numbers of detected genes were shown at the top of each box.

To determine the appropriate experimental conditions, several exploratory experiments were carried out on the E14 mouse embryonic stem cell line using the FACS workflow. Different RNase inhibitors and open chromatin tagging (ATAC) temperatures were tested. Key performance metrics from the experiments showed that incubation at 30 °C during open chromatin tagging provided the highest RNA-seq library quality: medians of over 17,000 total UMIs and more than 4,000 detected genes were observed in the 30 °C condition, and only over 2,000 total UMIs and around 1,000 detected genes were obtained in the 37 °C condition (**Supplementary Fig. 3a** and **b**). In addition, the performance of the two commonly used RNase inhibitors was similar, with the Protector RNase inhibitor being slightly better than the RiboLock (**Supplementary Fig. 3a** and **b**). The ATAC libraries generated by 30 °C tagmentation gave comparable quality to those generated by the commonly used 37 °C tagmentation condition, with similar number of unique nuclear fragments, fraction of reads in peaks (FRiP) and mitochondrial content (**Supplementary Fig. 3c-f**). Having determined the reaction conditions, species mixing experiments using human HEK293T and mouse NIH3T3 cells were performed to benchmark the method. Species assignment results were consistent between the two modalities (**Supplementary Fig. 3g** and **Methods**), with observed doublet rates of 4.2% and 8.8% for the FACS- and droplet-based workflow, respectively.

Next, to evaluate the feasibility of the method and the quality of the produced data, ISSAAC-seq was applied on several widely used cell lines using both FACS- and droplet-based strategies (**Supplementary Fig. 4a** and **b**). Furthermore, human peripheral blood mononuclear cells (PBMCs), a commonly used primary sample type, were also tested. For the droplet-based approach, we used the 10x Chromium Single Cell ATAC kit^19^ due to its wide availability in the community and modified some key steps to fit our workflow (see **Methods**). In both FACS- and droplet-based approaches, the ATAC-seq library insert fragments in ISSAAC-seq exhibited canonical nucleosomal ladder patterns that are observed in typical scATAC-seq experiments (**Supplementary Fig. 4b, c and e**). Reads from the ATAC-seq library were highly enriched around the transcription start site (TSS) (**Supplementary Fig. 4d** and **e**). The RNA-seq library fragments showed a unimodal distribution with a broad peak around 200 - 400 bp (**Supplementary Fig. 4b**), and a high proportion of reads from the RNA library came from exons (**Supplementary Fig. 4f**). In addition, there was significant amount of intronic reads in the RNA-seq library of ISSAAC-seq, which is commonly observed in single nucleus RNA-seq^36^. Visual inspection of the aggregated signals from single cells on a genome browser indicated ISSAAC-seq data were of high quality in different cell lines (**Supplementary Fig. 5a** and **b**). In K562 cells, ISSAAC-seq produced similar profiles compared to SHARE-seq, a recently developed open source single cell multi-omics method^7^, and the 10x Genomics Single Cell Multiome ATAC + Gene Expression kit (the 10x Multiome kit), a commercial multi-omics method (**Fig. 1b** and **Supplementary Fig. 5a**). When examining key performance metrics of ATAC and RNA libraries, we found ISSAAC-seq performed pretty well in both modalities, with over tens of thousands of unique nuclear reads in peaks (ATAC) and more than several thousand UMIs and detected genes (RNA) (**Fig. 1c** and **d, Supplementary Fig. 4g**). Within the same cell line or sample type, ISSAAC-seq generated data with comparable qualities to the commercial 10x Multiome kit, both of which had higher complexity (**Fig. 1c**) and sensitivity (**Fig. 1d**) than other similar methods. In addition, known cell types from PBMCs were successfully identified with ISSAAC-seq (**Supplementary Fig. 5c**). These observations suggested both the ATAC data and the RNA data from ISSAAC-seq were of high quality (**Supplementary Note 1**).

Having confirmed the feasibility of ISSAAC-seq, we next tested if the method was able to identify distinct cell types of a complex tissue. To this end, nuclei from the frozen adult mouse cerebral cortex were isolated and subjected to the ISSAAC-seq workflow, with both FACS- and droplet-based strategies (**Fig. 2a**). Similar to the results from the cell lines, we obtained medians of 16,464 and 16,619 unique reads in peaks from the ISSAAC-seq ATAC data by the FACS- and droplet-based strategy, respectively (**Fig. 2b**), which is comparable to the ATAC data from the 10x Multiome kit and better than all other open source methods (**Fig. 2b**). The RNA data of ISSAAC-seq were better than those of Paired-seq and SNARE-seq in terms of numbers of UMIs and detected genes (**Fig. 2b**). Compared to the results generated by the 10x Multiome kit and SHARE-seq, ISSAAC-seq RNA data detected similar amount of UMIs and genes (**Fig. 2b**). Then a computational pipeline was developed for exploring the cell types at both gene expression and chromatin accessibility layers (see **Methods** and **Supplementary Fig. 6a**), and a total of 10,378 nuclei were recovered and passed quality control (**Supplementary Fig. 6b**). The ISSAAC-seq RNA and ATAC data were visualized by uniform manifold approximation and projection (UMAP)^37^ on a two-dimensional space, separately (**Fig. 2c** and **d**). Unsupervised clustering identified 23 distinct RNA clusters and 22 different ATAC clusters (**Fig. 2c** and **Supplementary Fig. 7a**).

**Fig. 2.**
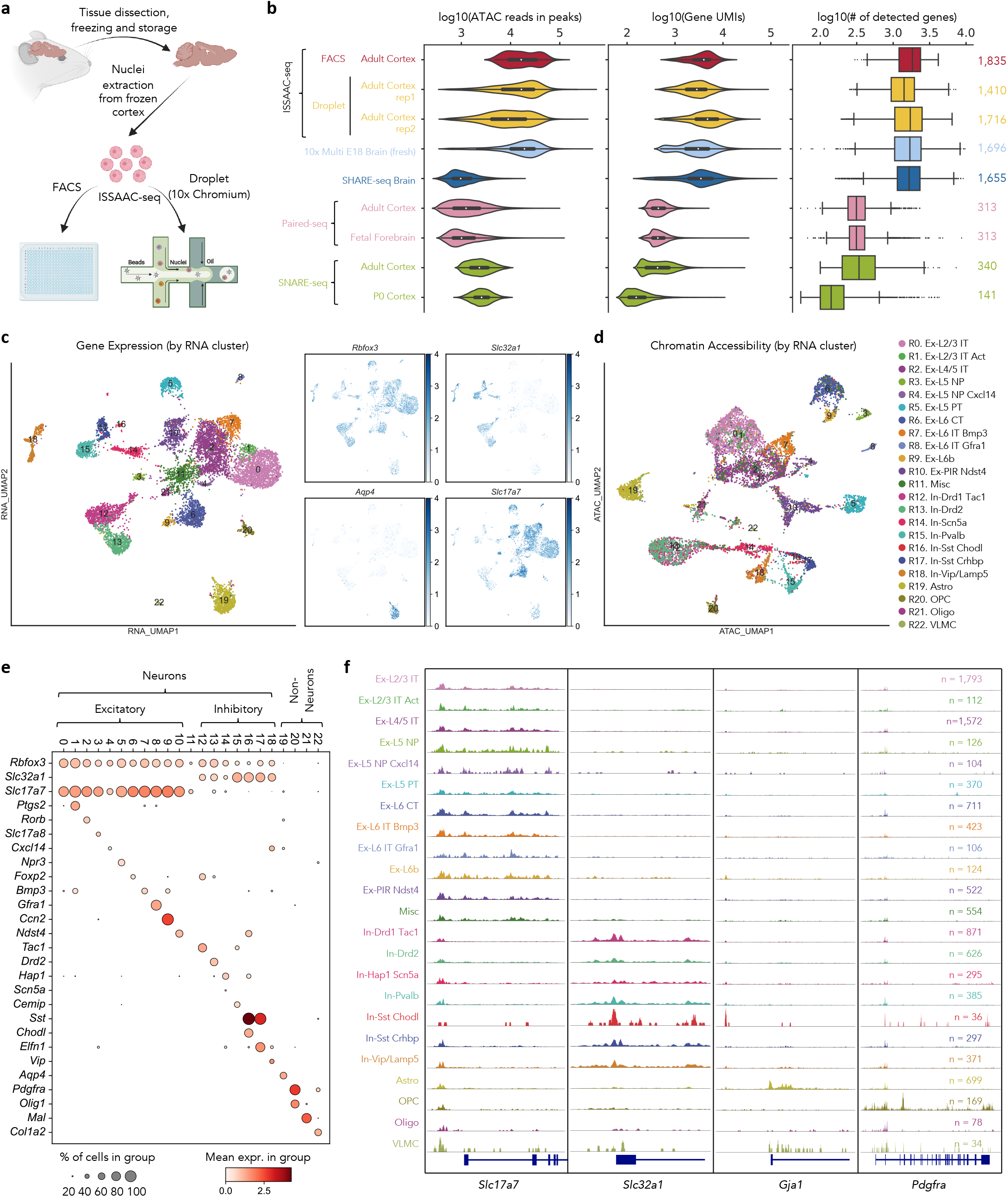
Simultaneous investigation of chromatin accessibility and gene expression from the mouse cerebral cortex using ISSAAC-seq. **a**, A schematic view of the experimental design for the mouse cerebral cortex experiments. **b**, Distributions of the numbers of reads in peaks (ATAC), UMIs (RNA) and detected genes (RNA) from different methods performed in the mouse brain samples. The medians of number of detected genes were indicated at the right hand side. **c**, UMAP projection of ISSAAC-seq RNA data coloured by the RNA clustering and expression values of indicated genes. **d**, UMAP projection of ISSAAC-seq ATAC data coloured by RNA clusters and cell types. **e**, Dotplot showing marker gene expression levels in each RNA cluster. **f**, UCSC genome browser tracks of aggregated scATAC-seq signals in each RNA cluster around four signature genes.

To figure out the cell types in the sample, we integrated the ISSAAC-seq RNA data with a mouse cortex scRNA-seq data set from the Allen Brain Institute^38^ using a label transfer technique^39^ to predict the cell type of each single cell in our data (**Supplementary Fig. 8** and **Supplementary Table 1**). Furthermore, we performed differential expression tests to find out the marker genes of each cell cluster. By combining these two approaches, we successfully identified twelve types of excitatory neurons (*Rbfox3+, Slc32al-, Slc17a7+*), seven types of inhibitory neurons (*Rbfox3+, Slc32a1+, Slcl7a7-*), and four types of non-neurons (*Rbfox3-*) (**Fig. 2c** and **e**, **Supplementary Fig. 9**). Although there appeared to be some minor difference between the two biological replicates in the In-*Scn5a* and Ex-PIR *Ndst4* neurons (see **Supplementary Note 2**), most cell clusters were robust and independent of batches (**Supplementary Fig. 7a** and **9**). Aggregates of the ATAC-seq signals in each cluster exhibit clear and specific open chromatin peaks around the marker gene loci (**Fig. 2f**).

Since each cell has matched gene expression and chromatin accessibility data, we could directly compare the RNA and ATAC clusters independent of integration methods. We labelled each cell on the ATAC UMAP space using RNA cluster label (**Fig. 2d**). In most cases, cells within the same RNA cluster also appeared in the same ATAC cluster (**Fig. 2d** and **Supplementary Fig. 7b**), indicating there is a general congruence between gene expression profile and chromatin accessibility. This is consistent with the results from recent single cell multi-omics studies^2,4,5,7^. Interestingly, we noticed that Ex-L4/5 IT neurons (RNA cluster R2) have three different chromatin accessibility states (ATAC clusters A2, A3 and A9), despite their similar gene expression profiles (**Fig. 3a** and **Supplementary Fig. 7 a** and **b**). If using the ATAC cluster labels as the guidance, we were able to detect some difference among the three ATAC clusters at the transcriptomic level (**Supplementary Fig. 7c-e**). It indicates that some heterogeneity at the gene expression level exists in the Ex-L4/5 IT neurons, but the difference is subtle and cannot be detected in the original clustering analysis with the RNA data only (**Fig. 3a**). This is consistent with previous reference atlas studies which showed that Ex-L4/5 IT neurons contained intermediate cells and the heterogeneity was not visible by clustering at the transcriptomic level^38,40^. In the ATAC data from ISSAAC-seq, it was apparent at the chromatin level that Ex-L4/5 neurons have at least three major subtypes (**Fig. 3a** and **Supplementary Fig. 7 a** and **b**). To further investigate the epigenetic differences among the three major ATAC clusters, cells from the same ATAC cluster were aggregated and peak calling was performed to identify open chromatin regions in each cluster. The top 1,000 cluster-specific peaks sorted by fold enrichment (see **Methods**) were taken for further analysis. Most cluster-specific peaks were located in the intergenic and intronic regions (**Fig. 3b**). When compared to the histone modifications occupancy data of the same types of neurons from the PairedTag data^26^, we found the cluster-specific peaks tend to locate in the regions marked by H3K4me1 and H3K27ac, indicating they might be potential enhancers (**Fig. 3c**). In addition, different transcription factor family motifs, such as T-box, basic helix-loop-helix (bHLH) and AP1, were enriched in the three sets of cluster-specific peaks, respectively (**Fig. 3d**), which suggests members from those families are important to establish the cluster specific chromatin states. SCENIC^41^ analysis of the gene expression data returned a few transcription factors in those families with higher activity specifically in one of the ATAC clusters (**Supplementary Fig. 7f**). The phenomenon that cells with similar gene expression profiles exhibit distinct chromatin accessibility status has been reported in mouse embryonic stem cells^42^, mouse T cells^43^ and human cortical neurons^44^. The functions of the chromatin difference remain to be explored, but the observation suggests those cells may have distinct chromatin potentials^7^ to react differently upon stimulation. These results demonstrated the direct comparison of ATAC and RNA profiles from the same cell helps uncover heterogeneity that will otherwise missed using only one modality.

**Fig. 3.**
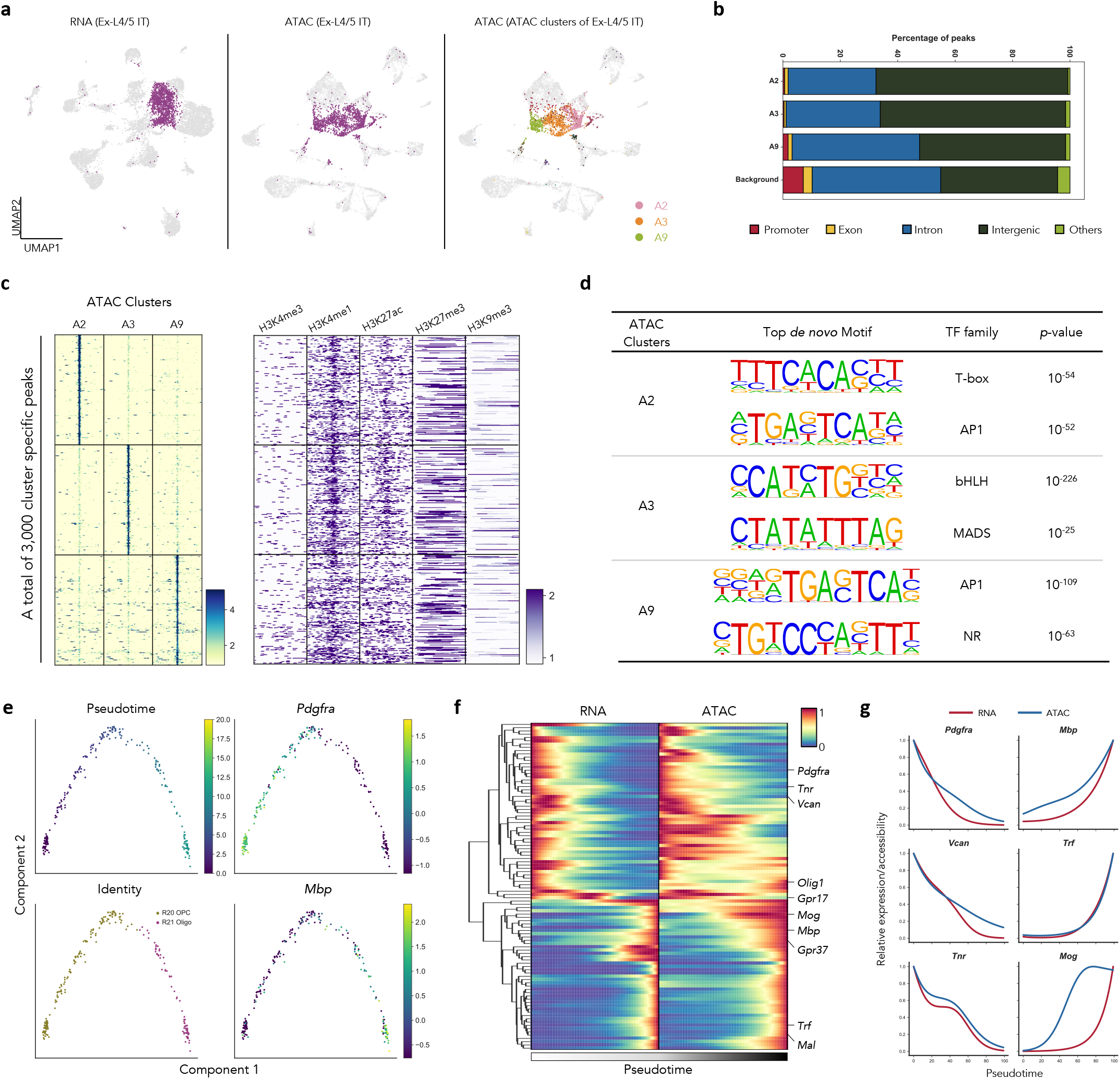
Joint analysis of RNA and ATAC from the same cell in the mouse cortex. **a**, UMAP projection of ISSAAC-seq RNA data (left) and ATAC data (middle and right). The RNA cluster R2 (Ex-L4/5 IT neurons) was highlighted by colours. All the rest cells were coloured by grey. **b**, Genomic distribution of the top 1000 cluster specific peaks and all ATAC peaks (background). **c**, Heatmap representation of the aggregated single cell ATAC-seq and PairedTag histone modification signals around the top 1000 cluster specific peaks in ATAC clusters A2, A3 and A9. **d**, The top *de novo* motifs enriched in the top 1000 cluster specific peaks of A2, A3 and A9. **e**, Pseudotime analysis of the oligodendrocyte progenitor cells (R20) and mature oligodendrocytes (R21). **f**, Heatmap representation of relative expression levels of genes that were differentially expressed along the pseudotime trajectory. The relative changes of chromatin accessibility in the same cell along the same trajectory were shown to the right. **g**, Examples of dynamic changes of gene expression and chromatin accessibility during the pseudotime.

The cell type annotation successfully identified two groups of cells in the oligodendrocyte lineage: oligodendrocyte progenitor cells (OPCs) and mature oligodendrocytes (Oligos) (**Fig. 2c**). A pseudotime analysis^45^ was carried out on those cells to investigate the oligodendrocyte maturation process. Since most pseudotime inference tools are designed to work with gene expression data, top differentially expressed genes between OPCs and Oligos were used to arrange cells in a pseudotime trajectory (**Fig. 3e**). Known markers exhibit expected dynamics along the trajectory, indicating the pseudotime indeed represents the oligodendrocyte maturation process. For examples, expression levels of the OPC marker *Pdgfra* and the mature Oligo marker *Mbp* went down and up along the trajectory, respectively (**Fig. 3e**). Next, taking the advantages of our multi-omics data, we checked the gene activity score measured by chromatin accessibility from the same cell along the same oligodendrocyte maturation trajectory (**Fig. 3f**). Distinct patterns of gene expression and chromatin accessibility became prominent in this analysis. In general, changes in gene expression levels were concordant with dynamics of chromatin accessibility in the trajectory (**Fig. 3f** and **g**). Genes with increasing expression levels along the trajectory also exhibited increasing chromatin accessibility, but the increase at the chromatin level preceded the gene expression change in many genes, such as *Mbp* and *Mog* (**Fig. 3f** and **g**). This is consistent with a recent finding that chromatin become accessible prior to the start of the corresponding gene expression during lineage commitment^7^. Interestingly, the expression and chromatin dynamics of genes with decreasing levels along the trajectory showed a different pattern. Although both gene expression and chromatin accessibility were getting lower, changes of chromatin accessibility were slower or lagging behind of gene expression (**Fig. 3f** and **g**). This observation indicated the change of chromatin status may lead to the silence of gene expression, while the fully close of promoters need longer time. These results demonstrated that ISSAAC-seq could be used to investigate the gene regulatory mechanisms during dynamic processes, such as cellular differentiation.

In general, our method provides a robust and highly sensitive way of investigating gene expression and chromatin accessibility from the same cell. Compared to the commercial 10x Multiome kit, ISSAAC-seq offers data with comparable qualities at a much lower cost (**Supplementary Note 3**). The method has a very flexible workflow that can be combined with either FACS or any droplet systems with a Nextera capture sequence, such as 10x Genomics^19^, Bio-Rad^20^ and HyDrop^46^. The FACS workflow is suitable for limited cell number, and the droplet workflow can be used for high-throughput cell profiling. The library structures generated by ISSAAC-seq is similar to the standard Illumina library (**Supplementary Figs. 1** and **2**), which does not require custom sequencing primers or recipes. Furthermore, with the recent development of multi-omics methods that utilize antibody-oligo conjugates, such as TEA-seq^28^, DOGMA-seq^27^ and NEAT-seq^47^, it is possible to capture ATAC, RNA and protein information from the same cell. Similar design can also be applied in the ISSAAC-seq method to collect additional information (see **Supplementary Note 4**). The procedures are also modular, containing an *in situ* reaction module, a cell isolation module and a library construction module. The latter two modules have almost the same steps in many existing single cell methods. Therefore, our method serves as a valuable tool for single cell multi-omics profiling and can be adapted straightforwardly in many labs.

## Figure Legends

**Supplementary Fig. 1.**
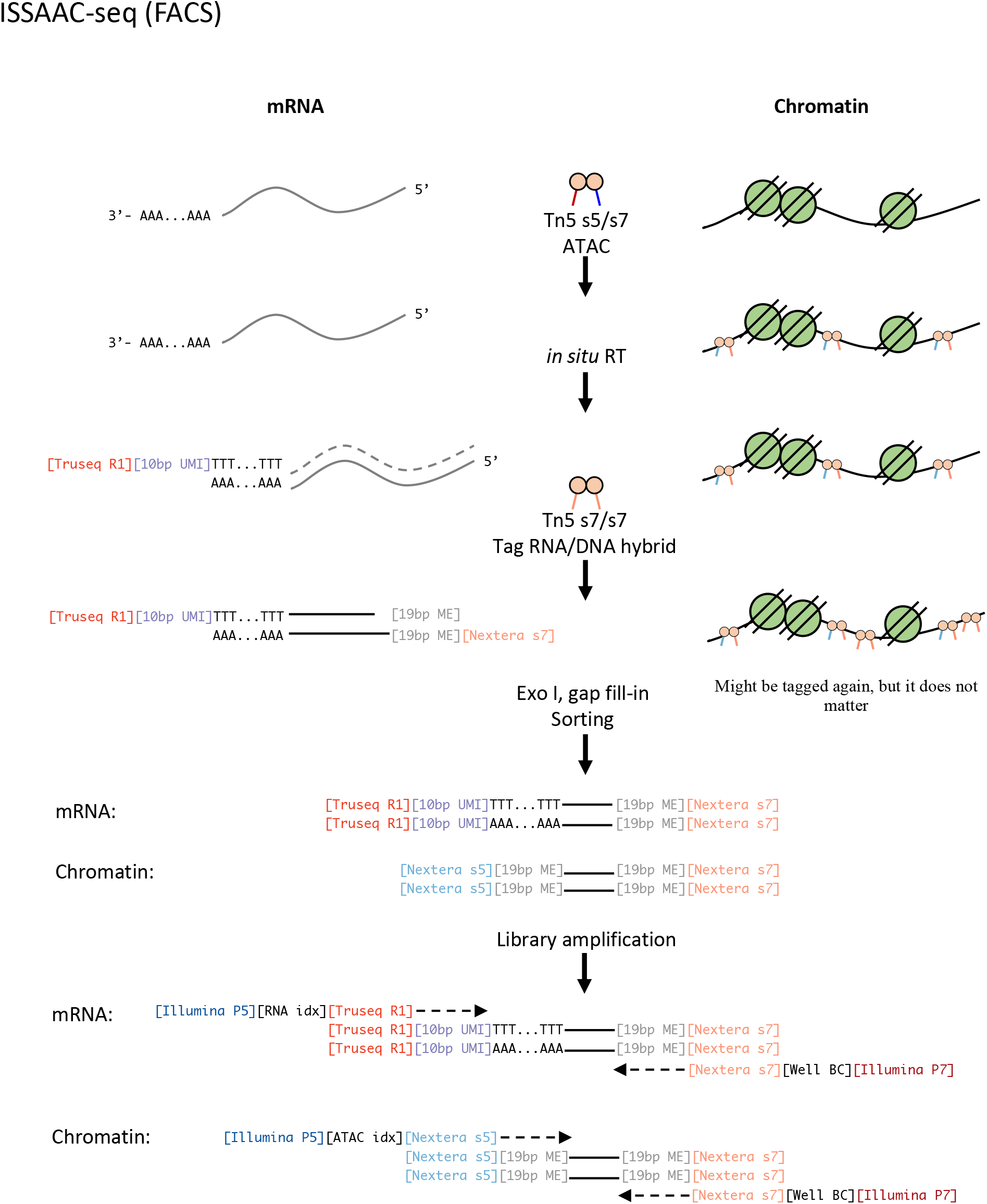
Schematic view of the plate-based (FACS) ISSAAC-seq workflow and adaptor configurations.

**Supplementary Fig. 2.**
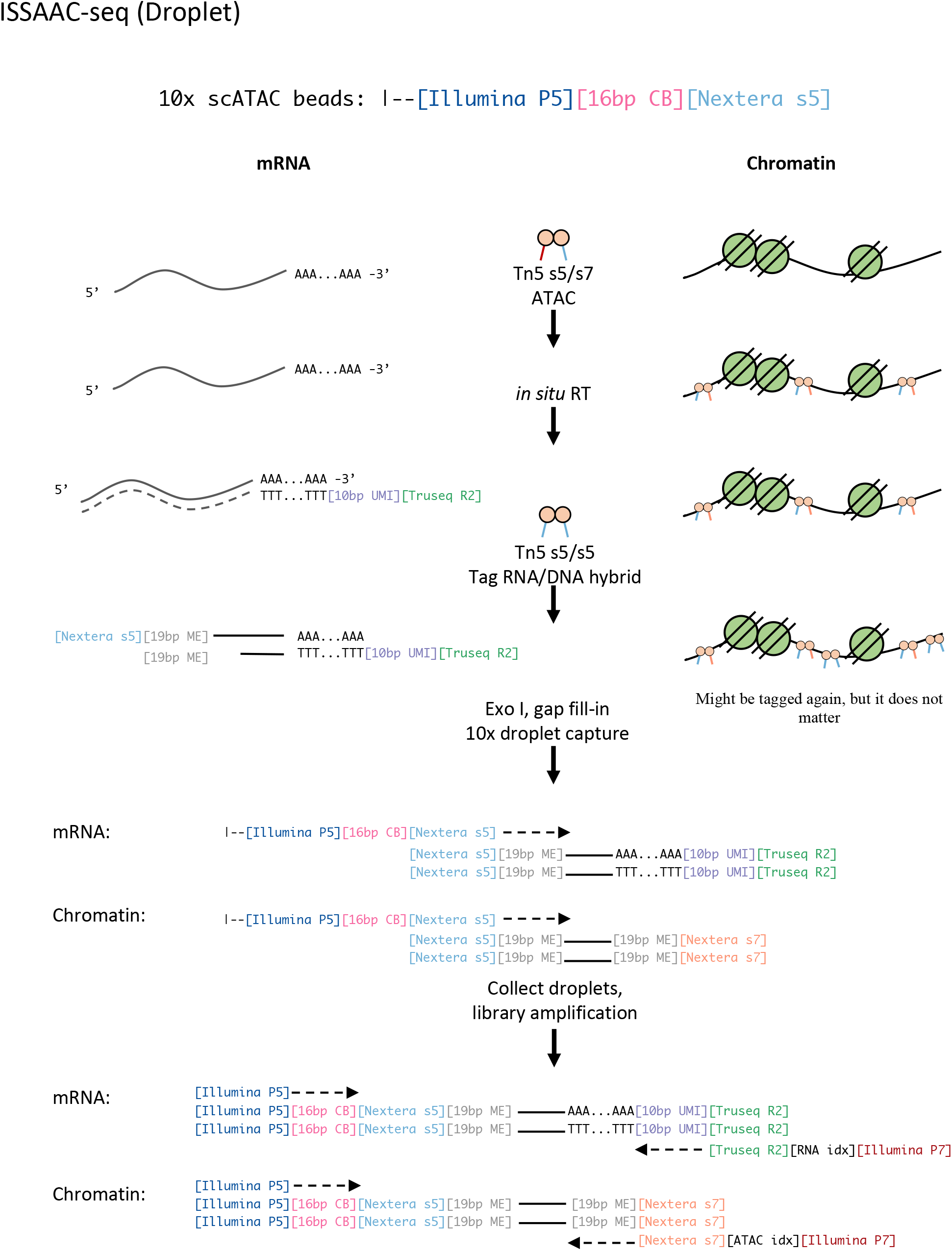
Schematic view of the droplet-based ISSAAC-seq workflow and adaptor configurations.

**Supplementary Fig. 3.**
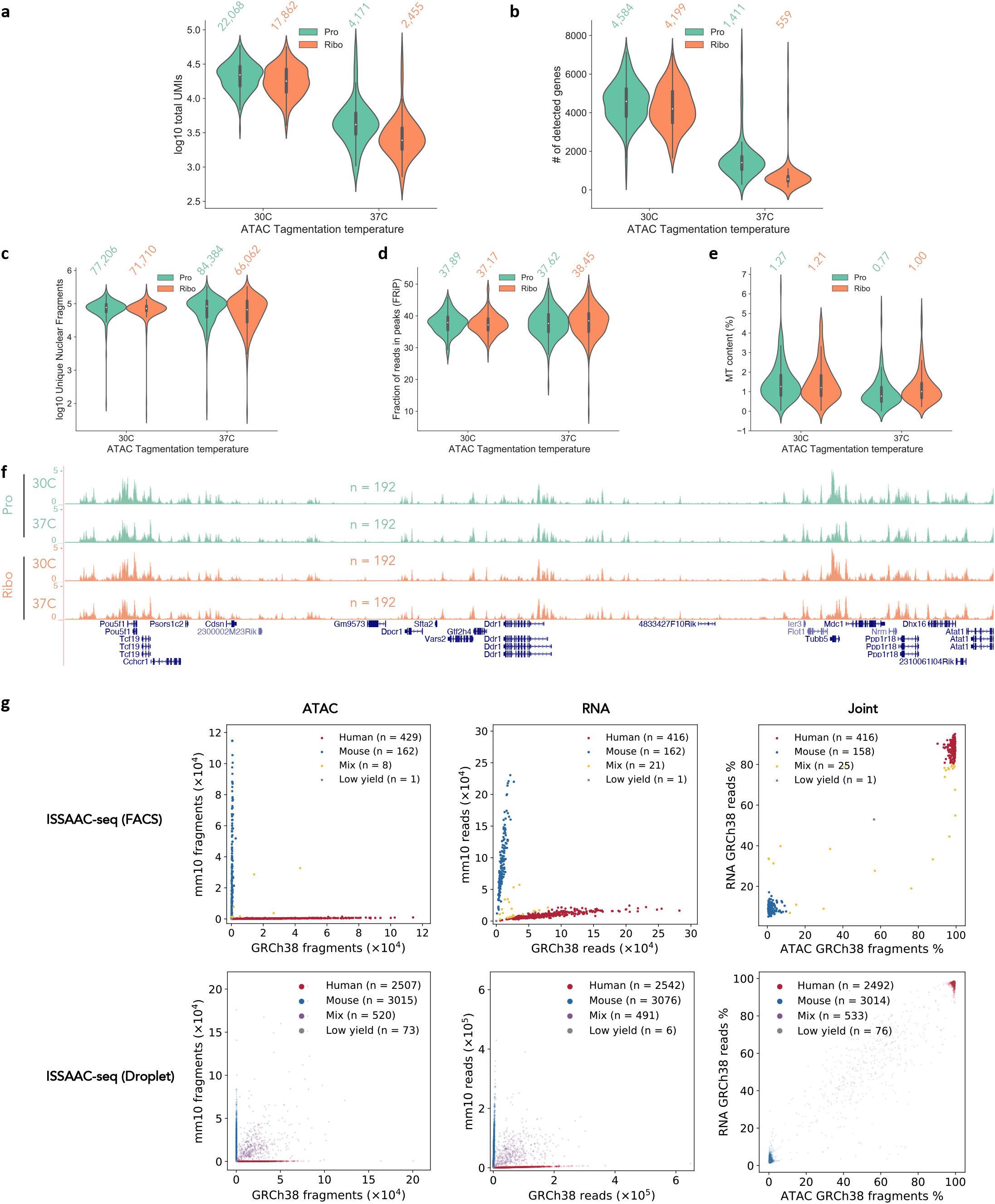
Experimental tests and benchmarking of ISSAAC-seq. **a** and **b**, Assessment of ISSAAC-seq RNA data. Distributions of total UMIs (**a**) and numbers of detected genes (**b**) from indicated conditions performed on the E14 mouse embryonic stem cells. Pro, Protector RNase inhibitor. Ribo, RiboLock RNase inhibitors. **c**-**e**, Assessment of ISSAAC-seq ATAC data. Distributions of unique nuclear fragments (**c**), fraction of reads in peaks (FRiP) (**d**) and mitochondrial content (**e**) from indicated conditions performed on the E14 mouse embryonic stem cells. **f**, UCSC genome browser tracks of aggregated scATAC-seq signal from indicated conditions. **g**, Species mixing experiments of ISSAAC-seq using HEK293T and NIH3T3 cells.

**Supplementary Fig. 4.**
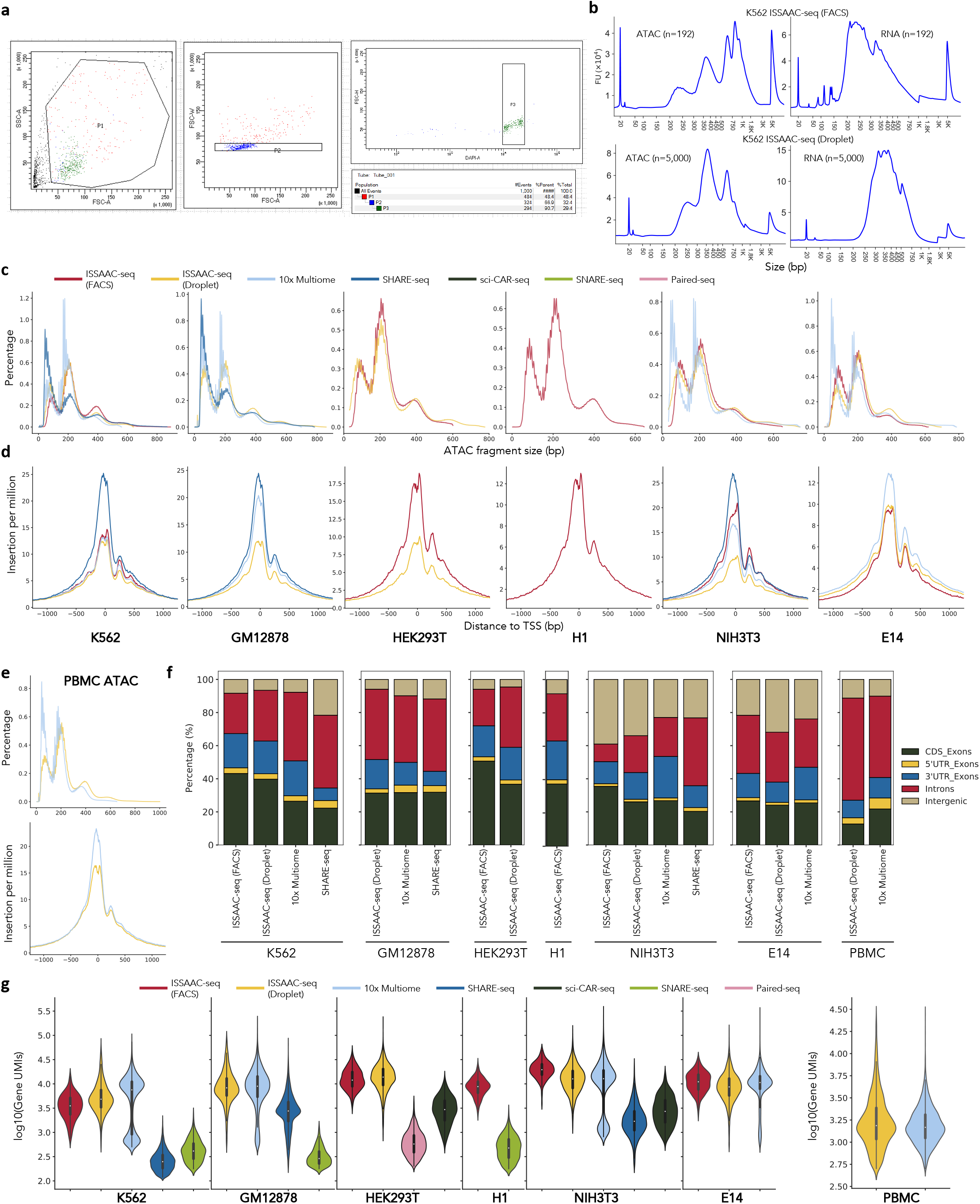
ISSAAC-seq experimental results from different cell lines and PBMCs. **a**, Examples of a typical FACS based workflow using K562 cells. FSC-A and SSC-A were used to select intact nuclei (P1). FSC-A and FSC-W were used to select singlets (P2). DAPI-A was used to select DAPI positive nuclei (P3). **b**, DNA electrophoresis of typical ISSAAC-seq libraries from the FACS (left) and droplet (right) workflow using K562 cells. **c** and **d**, Fragment size (**c**) and Tn5 insertion frequency (**d**) of ATAC data using different methods in the indicated cell lines. **e**, quality metrics of the ISSAAC-seq PBMC data. **f**, Genomic distribution of reads from RNA data using different methods. **g**, Distributions of the numbers of UMIs using different methods performed in the indicated cell lines and PBMCs.

**Supplementary Fig. 5.**
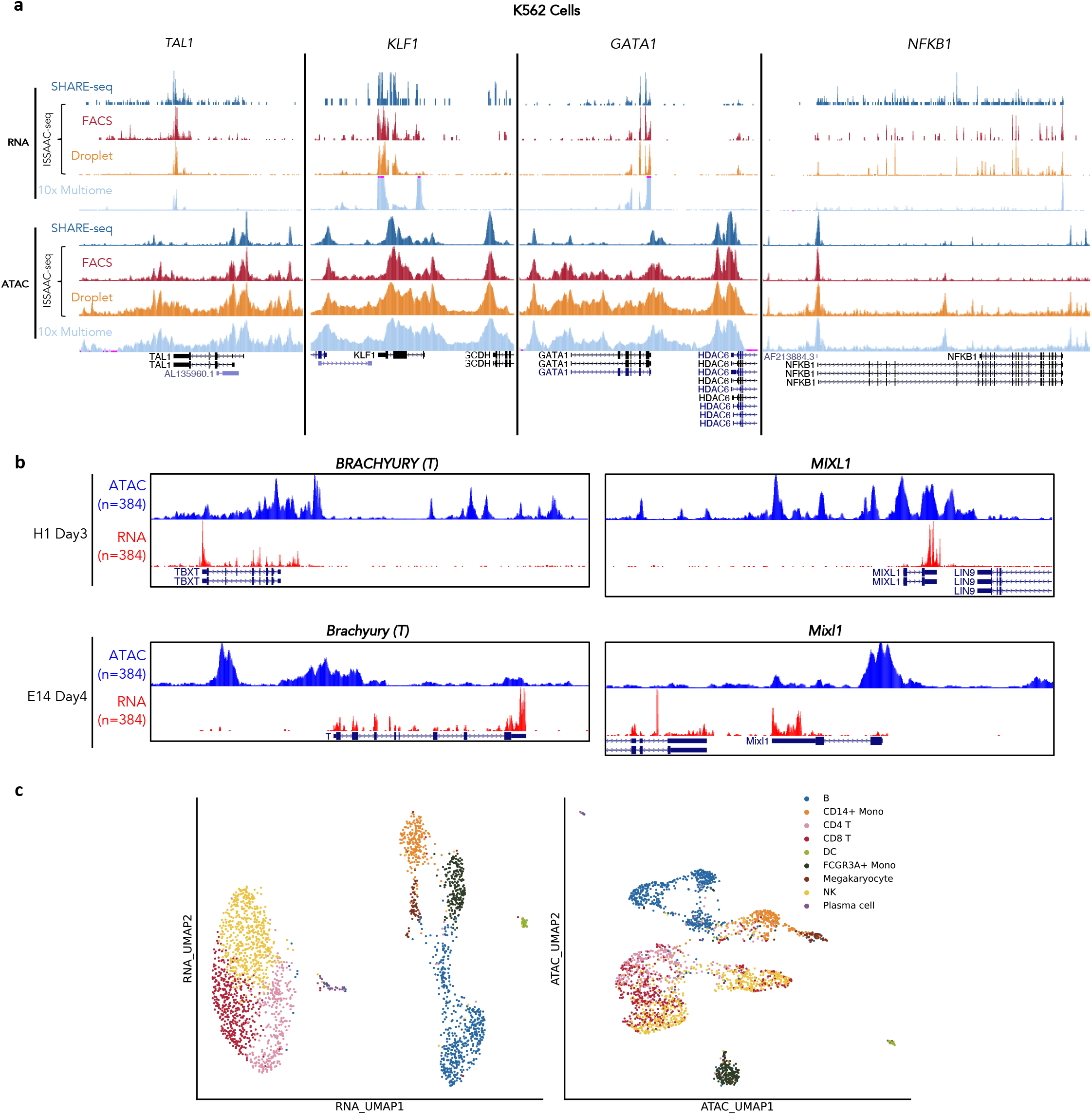
UCSC genome browser tracks of single cell aggregates from ISSAAC-seq. **a**, Visual comparisons of SHARE-seq, ISSAAC-seq and the 10x Multiome kit from K562 cells. Cell numbers are the same as in **Fig. 1b**. **b**, Visualisation of ATAC and RNA signals from single cell aggregates of ISSAAC-seq (FACS) in H1 (Day 3 upon mesoderm differentiation) and E14 (Day 4 upon EB differentiation) cells around the *BRACHYURY* and *MIXL1* genes. **c**, UMAP projection of ISSAAC-seq PBMCs.

**Supplementary Fig. 6.**
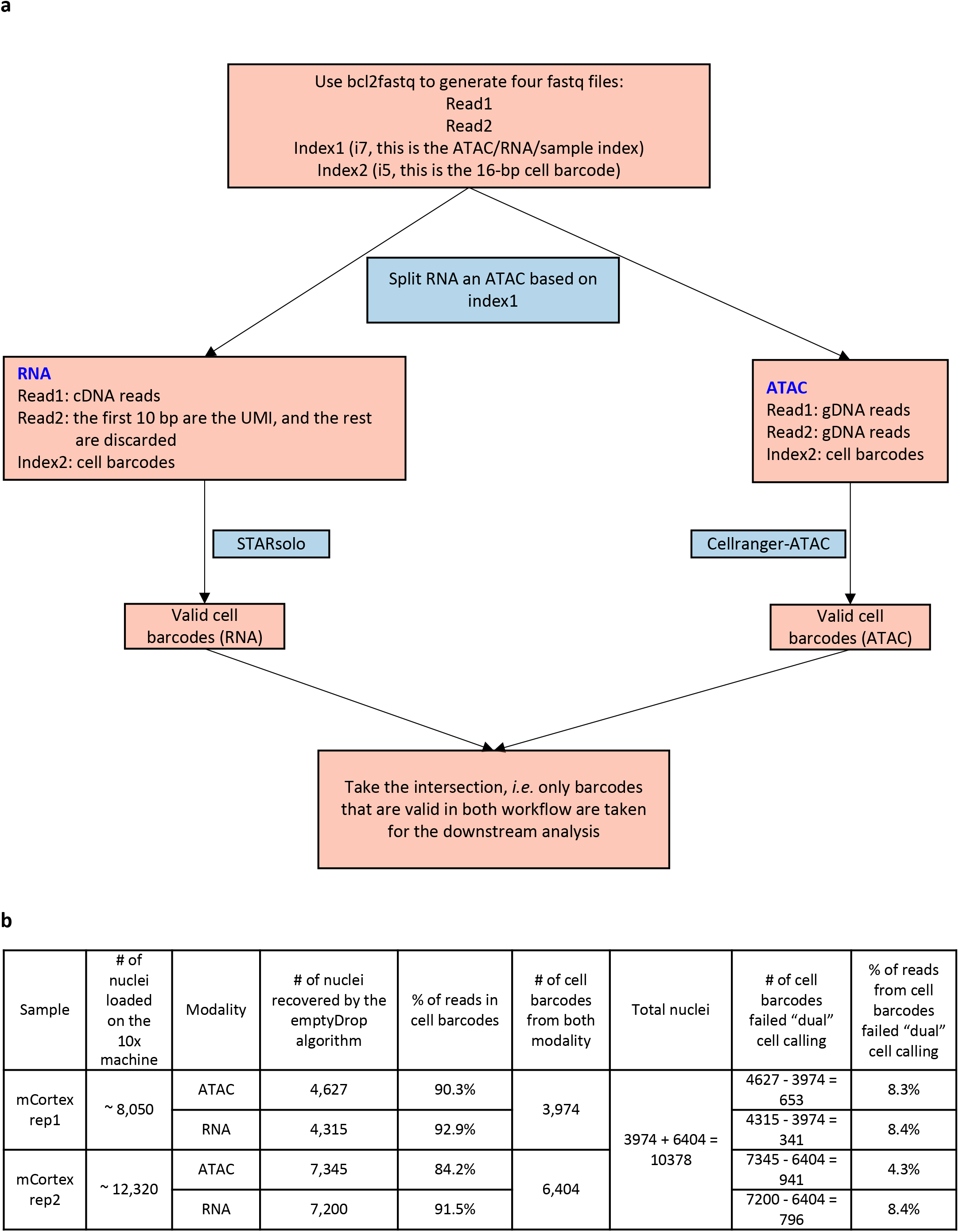
Computational pipeline for the data processing of droplet-based ISSAAC-seq. **a**, Flow chart of the computational pipeline to process mouse cortex data. **b**, Cell barcode qualities and the number of nuclei returned by the pipeline in each step.

**Supplementary Fig. 7.**
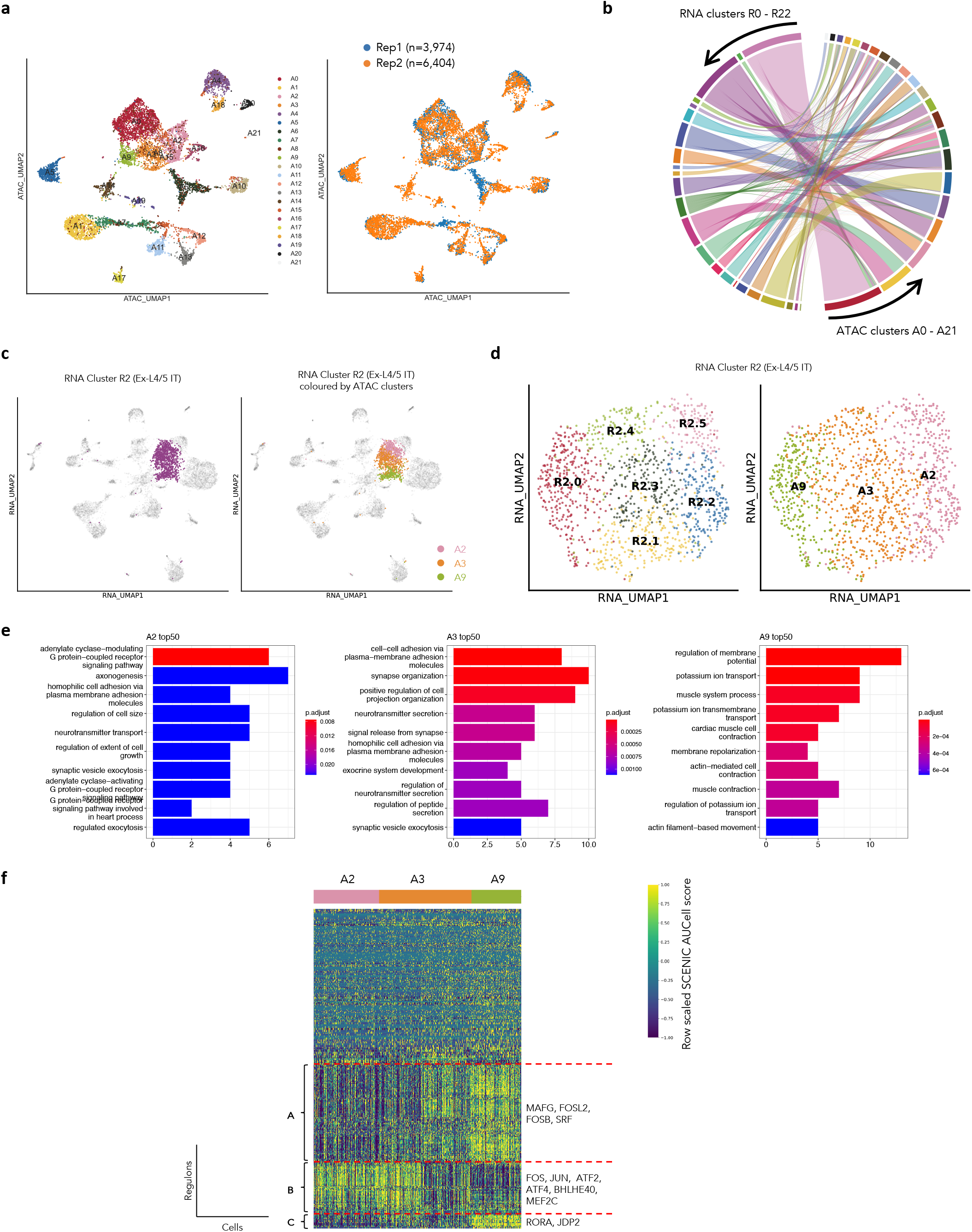
ISSAAC-seq ATAC and RNA data comparison. **a**, UMAP projection of mouse cortex cells with ISSAAC-seq ATAC data coloured by ATAC clusters (left) and batches (right). **b**, Ribbon plot showed the relationship between all RNA clusters and all ATAC clusters. **c**, UMAP projection of mouse cortex cells with ISSAAC-seq RNA data, highlighted by RNA cluster R2 Ex-L4/5 IT neurons (left) and ATAC cluster A2, A3 and A9 (right). **d**, UMAP projection of Ex-L4/5 IT neurons (RNA cluster R2) coloured by RNA subclusters (left) and ATAC clusters. **e**, Gene ontology of the top 50 differentially expressed genes among A2, A3 and A9 clusters within the Ex-L4/5 IT neurons. **f**, SCENIC analysis of regulon (TF) activity in A2, A3, and A9 cells.

**Supplementary Fig. 8.**
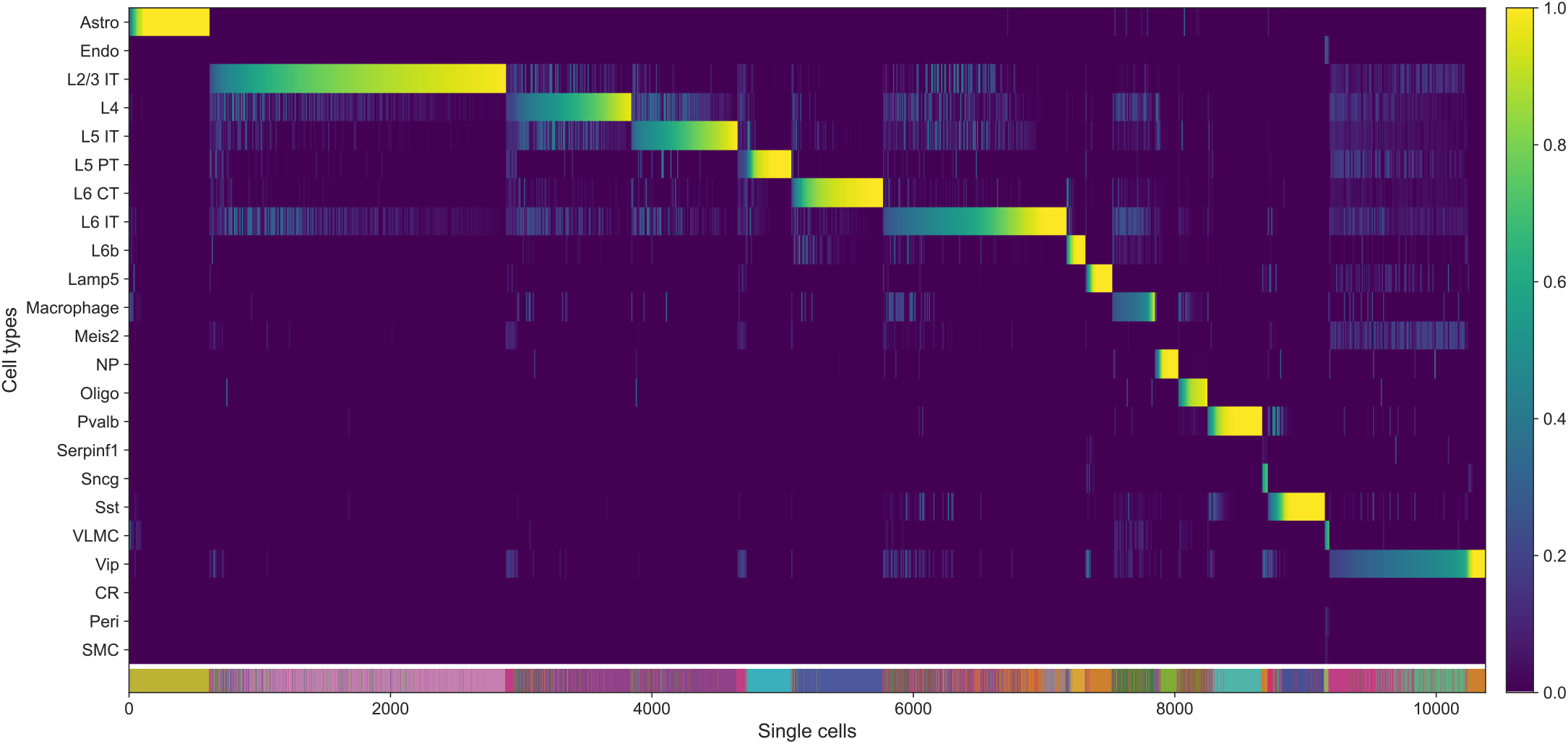
The prediction probability of each single cell based on the Allen Brain reference data using the label transfer function from Seurat.

**Supplementary Fig. 9.**
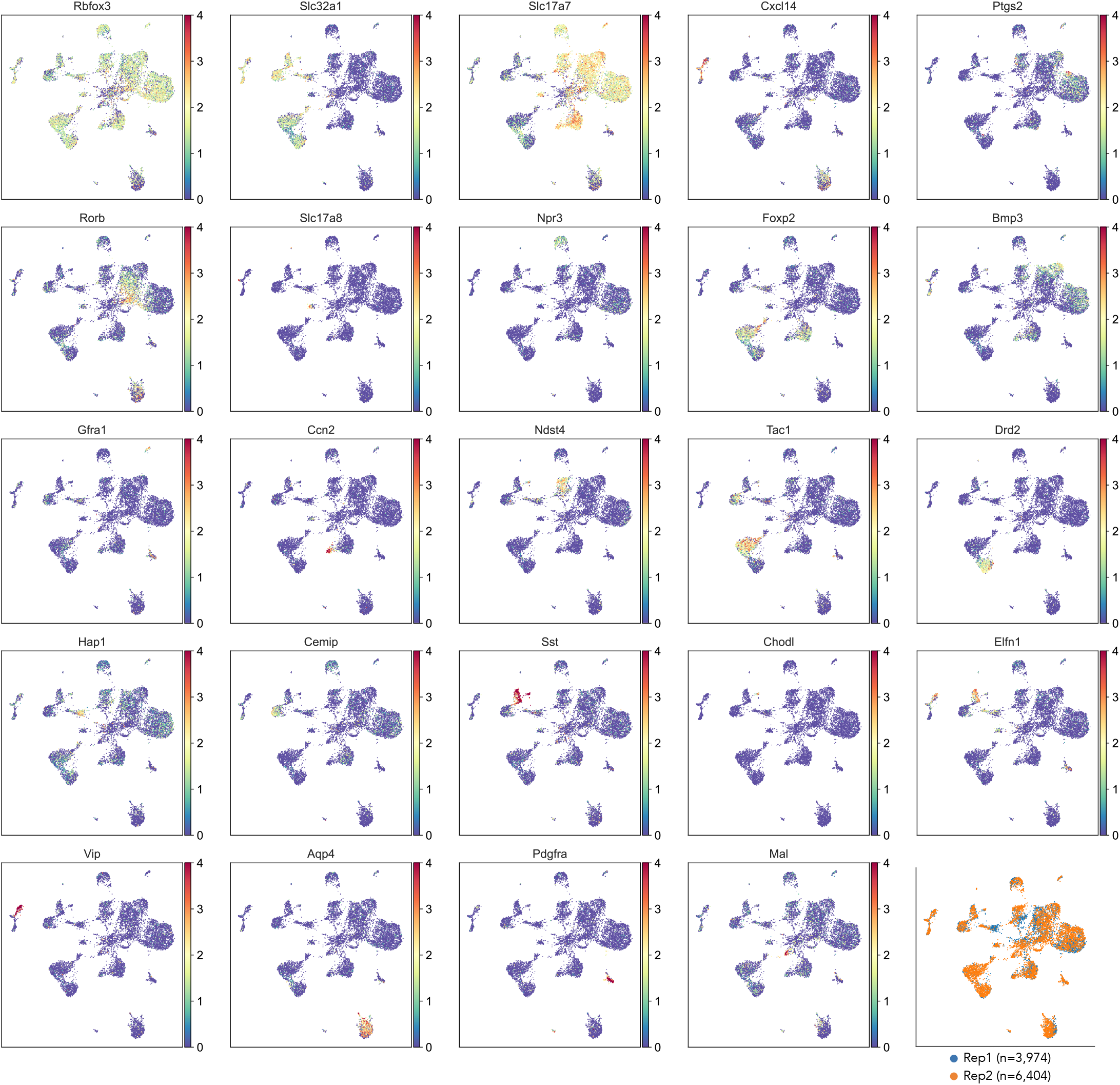
UMAP projection of ISSAAC-seq RNA data coloured by the expression levels of indicated genes or batches.

## Author contributions

X.C and W.J conceived the project. X.C, W.X and W.J designed the protocol and experiments. W.X performed the experiments with help from Y.C, N.H, Q.Z, X.W and Y.H. W.Y carried out the computational analysis with help from Y.Z, W.X, X.C, W.J and K.S. X.C, W.J, K.S and N.H supervised the entire project. All authors contributed to the writing.

## Acknowledgement

We thank all members from the Chen and Jin labs for the helpful discussion of the project. We also thank Prof. Wei Chen and Dr. Huanhuan Cui for the helpful advice on the experiments. We thank Xibin Lu for the excellent support of FACS. We acknowledge the assistance of SUSTech Core Research Facilities. The computational work was supported by Center for Computational Science and Engineering at Southern University of Science and Technology. Figure 1a and Figure 2a were created with BioRender.com.

## Methods

### Cell culture

Cells were cultured according to standard procedures in RPMI 1640 (Gibco, cat. no. 21870076) for K562 and GM12878 cells or DMEM (Hyclone, cat. no. SH30243.01) for HEK293T and NIH3T3 cells, supplemented with 10% FBS (Hyclone, cat. no. 30160), 100 U/ml penicillin and 100 μg/ml streptomycin sulfate (Hyclone, cat. no. SV30010). HEK293T and NIH3T3 cells were digested with 0.25% Trypsin (Thermo Fisher, cat. no. 25200056) for preparing single-cell suspension.

mESC E14 cells were cultured in DMEM supplemented with 10% embryonic stem cell qualified FBS (Gibco, cat. no. 30044333), 1000 U/ml mLif (Millipore, cat. no. ESG1107), 0.1 mM β-mercaptoethanol (Gibco, cat. no. 21985023), 100 U/ml penicillin and 100 μg/ml streptomycin sulfate on gelatin (Amresco, cat. no. 9764-100G) coated dishes. mESCs at 80% confluence were harvested using 0.05% Trypsin for preparing single-cell suspension. For embryoid body formation, 2×10^6^ undifferentiated mESCs were seeded onto bacteriological dishes containing 15 ml of ES medium without LIF. After 24 hours, the primary aggregates were spun down and transferred to new bacteriological dishes at the split ratio of 1:10 for further differentiation. The medium was changed every other day.

hESC H1 cells were cultured according to standard procedures in mTeSR1 (STEMCELL, cat. no. 05825) on Corning Matrigel hESC-Qualified Matrix (Corning, cat. no. 354277). For mesoderm differentiation, hESCs were differentiated using the STEMdiff Definitive mesoderm Kit (STEMCELL, cat. no. 05220) according to the manufacturer’s instructions. Briefly, hESCs at 80% confluence were harvested using StemPro Accutase cell dissociation reagent (ThermoFisher, cat. no. A1110501) and reseeded in a single-cell manner on Matrigel plates. Cells were incubated with STEMdiff™ Mesoderm Induction Medium for indicated time with daily medium replacement.

### Mice

Mice were maintained in laboratory animal center at Southern University of Science and Technology. Procedures were approved by Experimental Animal Welfare Ethics Committee, Southern University of Science and Technology. Normal brain cortex was collected from wild-type C57BL/6 mice aged 8-9 weeks.

### Nuclei preparation from cell line

Cells were harvested, washed once with ice-cold DPBS-0.5% BSA containing 0.2 U/μl RiboLock Rnase inhibitor (Thermo Fisher, cat. no. EO0382), counted and viability detected by Trypan blue (Thermo Fisher, cat. no.15250061). Cell viability should be above 90%. For each sample, 1×10^5^ cells were collected by centrifuge at 500 g, 4 °C, 5 min. The cells were resuspended in 50 μl of ice-cold RSB-DTN-RI (10 mM Tris-HCl, pH 7.4, 10 mM sodium chloride, 3 mM magnesium chloride, 0.1% Tween-20 (Sigma, cat. no. 655205-250ML), 0.1% IGEPAL CA-630 (Sigma, cat. no. I8896), 0.01% Digitonin (Promega, cat. no. G9441), 0.8 U/μl RiboLock RNase inhibitor) and incubated on ice for 3 min. The lysis was washed with 1 ml ice-cold RSB-T (10 mM Tris-HCl, pH 7.4, 10 mM sodium chloride, 3 mM magnesium chloride, 0.1% Tween-20) and nuclei were spun down at 1000g, 4°C, 8 min. All centrifugations in sample processing were performed on the swing bucket centrifuge.

### Nuclei isolation from mouse cortex

An adult mouse brain cortex was dissected, snap-frozen in liquid nitrogen, and stored at −80 °C. For single nucleus isolation, frozen cortex (2-3 mm^3^) was placed into a pre-chilled 1 mL Dounce homogenizer with 1 mL of homogenization buffer (250 mM sucrose, 25 mM potassium chloride, 5 mM magnesium chloride, 10 mM Tris-HCl pH 8.0, 1 μM DTT, 1x Protease Inhibitor Cocktail (Roche, cat. no. 11697498001), 0.4U/μl RNasin (Promega, cat. no. N2111), and 0.1% (v/v) TritonX-100). Tissue was homogenized with 5 strokes of the loose pestle, followed by 10 strokes of the tight pestle. The sample was centrifuged at 100 g, 4 °C for 1 min to remove large debris. The supernatant was filtered through a pre-chilled 2 mL round bottom tube with cell-strainer cap (Falcon), and the effluent was transferred to a pre-chilled 1.5 ml tube, centrifuged at 1000 g, 4 °C for 3 min. The pellet was resuspended in 1 ml of ice-cold NIM2 buffer (250 mM sucrose, 25 mM potassium chloride, 5 mM magnesium chloride, 10 mM Tris-HCl pH 8.0, 1 μM DTT, 1x Roche Protease Inhibitor Cocktail, and 0.4 U/μl RNasin), and centrifuged at 1000 g, 4 °C for 3 min. Nuclei were then resuspended in 1 ml of ice-cold PBSI (1% (w/v) BSA, 1 μM DTT, and 0.4 U/μl RNasin).

### PBMC isolation

PBMCs from a healthy donor were isolated using Lymphoprep^™^ (STEMCELL, cat. no. 07801). Following isolation, PBMCs were subjected to RBC lysis, washing, and counting. PBMC aliquots were cryopreserved in 90% FBS+ 10% DMSO and stored in liquid nitrogen. For cell thawing, cryopreserved PBMCs were removed from liquid nitrogen storage and thawed in a 37 °C water bath for 3-5 min until no ice was visible. Cells were then washed once with 10 ml DPBS-0.5% w/v BSA. Dead cells are removed using Dead Cell Removal Kit (Miltenyi Biotec, cat. no. 130-090-101). 1×10^6^ PBMCs were crosslinked in 1 ml freshly prepared glyoxal fixation buffer (3% glyoxal, 0.75% glacial acetic acid, pH 5.0) at room temperature for 7 min, washed twice with 1 ml DPBS-0.5% w/v BSA, then hold on ice.

### Oligonucleotide sequences

The sequences of oligos used in this study and the purification methods can be found in **Supplementary Table 2**.

### Transposome complex assembly

To prepare the primers for ISSAAC-seq, ME_S5, ME_S7 or ME_bottom oligos was dissolved in annealing buffer (10 mM Tris-HCl, pH 8.0, 50 mM sodium chloride, 1 mM EDTA, pH 8.0) to a final concentration of 100 μM. 25 μl ME_S5 oligo (100 μM) was mixed with 25 μl ME_Bottom (100 μM), and annealed in a thermocycler as follows: 98 °C for 3 min, and slowly cooled to 16 °C with a temperature ramp of −0.1 °C/s, to generate S5_adaptor (50 μM). Similarly, ME_S7 and ME_Bottom oligos were annealed to form S7_adaptor.

To prepare regular Tn5-S5/S7 transposome complex, 12 μl S5_adaptor (20 μM), 12 μl S7_adaptor (20 μM), 48 μl purified Tn5 (0.5 μg/μl, Fapon Biotech, cat. no. NK001), and 88 μl coupling buffer (50 mM Tris-HCl, pH 7.5, 100 mM NaCl, 0.1 mM EDTA, pH 8.0, 0.1% Triton X-100, 1 mM DTT, 50% glycerol) were mixed and incubated at room temperature for 1 h. Tn5-S7/S7 homodimer transposome was assembled by mixing 24 μl S7_adaptor (10 μM), 48 μl purified Tn5 (0.5 μg/μl), and 88 μl coupling buffer. Similarly, Tn5-S5/S5 homodimer transposome was assembled by mixing 24 μl S5_adaptor (10 μM), 48 μl purified Tn5 (0.5 μg/μl), and 88 μl coupling buffer.

More details of the Tn5 assembly and quality assessment can be found in the protocols.io page: https://www.protocols.io/private/D62649C3B25011ECAC280A58A9FEAC02

### Chromatin tagmentation

For Chromatin tagmentation, nuclei were resuspended in 50 μl of Tn5-S5/S7 tagmentation mix containing 33 mM Tris-acetate, pH 7.8, 66 mM potassium acetate, 10 mM magnesium acetate, 16% dimethylformamide (DMF, Sigma, cat. no. D4551-250ML), 0.01% digitonin, 1.2 U/μl RiboLock Rnase inhibitor (Thermo Fisher, cat.no. EO0382), 0.4 U/μl SUPERaseIn (Thermo Fisher, cat.no. AM2694), 0.8 U/μl RnaseOUT (Thermo Fisher, cat. no. 10777019), and 5 μl Tn5-S5/S7 transposome complex. The tagmentation reaction was done on a thermomixer at 800 rpm, 30 °C, 30 min. The reaction was then stopped by adding equal volume (50 μl) of tagmentation stop buffer (10 mM Tris-HCl pH 7.8, 20 mM EDTA, pH 8.0, 2% BSA).

### *in situ* reverse transcription

After chromatin tagmentation, nuclei were spun down at 1000 g, 4 °C, 3 min, and washed twice with 200 μl of ice-cold 0.5× DPBS-0.5% BSA containing 0.2 U/μl RiboLock RNase inhibitor. Nuclei were resuspended in 100 μl of reverse transcription mix containing 0.5 mM dNTP, 10 U/μl Maxima H minus reverse transcriptase (Thermo Fisher, cat.no. EP0753), 0.8 U/μl RiboLock Rnase inhibitor, 0.2 U/μl SUPERaseIn, 0.4 U/μl RnaseOUT, 12% PEG8000, 50 mM Tris-HCl, pH 8.0, 75 mM sodium chloride, 3 mM magnesium chloride, 10 mM dithiothreitol, and 2 μM TruseqR1_oligo_dT for the plate-based workflow or 2 μM TruseqR2_oligo_dT for the 10x droplet workflow. The *in situ* reverse transcription reaction was performed as follows: 50°C 10 min; then 3 cycles of: 8°C 12s, 15°C 45s, 20°C 45s, 30°C 30s, 42°C 2 min, 50°C 3 min; followed by a final step at 50°C for 5 min.

### RNA/DNA hybrid tagmentation

After reverse transcription, nuclei were spun down at 800g, 4°C, 5 min, and washed twice with 200 μl of ice-cold 0.5× DPBS-0.5% BSA, then resuspended in 50 μl of tagmentation mix containing 33 mM Tris-acetate, pH 7.8, 66 mM potassium acetate, 10 mM magnesium acetate, 16% dimethylformamide (DMF, Sigma, cat. no. D4551-250ML), 0.01% digitonin, and 1.5 μl of Tn5-S7/S7 transposome for the plate-based workflow or 1.5 μl of Tn5-S5/S5 for the 10x droplet workflow. The tagmentation reaction was done on a thermomixer at 800 rpm, 37 °C, 30 min. The reaction was then stopped by adding equal volume (50 μl) of tagmentation stop buffer (10 mM Tris-HCl pH 7.8, 20 mM EDTA, pH 8.0, 2% BSA).

### EXO I digestion and gap fill-in

After RNA/DNA hybrid tagmentation, nuclei were spun down at 800 g, 4 °C, 5 min, and washed twice with 200 μl of ice-cold 0.5× DPBS-0.5%BSA. Then nuclei were resuspended in 50 μl of reaction mix containing 0.5 mM dNTP, 8 U/μl Maxima H minus reverse transcriptase, 2 U/μl Thermolabile Exonuclease I (NEB, cat. no. M0568), 50 mM Tris-HCl, pH 8.0, 75 mM sodium chloride, 3 mM magnesium chloride, and 10 mM dithiothreitol. EXO I digestion and gap fill-in reaction was done on a thermomixer at 800 rpm, 37 °C, 15 min.

### Single nuclei sorting and library pre-amplification in plate-based workflow

After EXO I digestion and gap fill-in, nuclei were resuspended in 400 μl of 0.5× DPBS-0.5% BSA and transferred to a FACS tube. DAPI (Thermo Fisher, cat. no. 62248) was added at a final concentration of 1 μg/μl to stain the nuclei. DAPI positive single nuclei were sorted into each well in a 384-well plate containing 3 μl lysis buffer (50 mM Tris-HCl, pH 8.0, 50 mM sodium chloride, 0.2% SDS, 10 μM N7xx primer, 10 μM Truseq_S5_short primer, 10 μM Nextra_S5_short primer) by FACS as previously described^34^. The plates were incubated at 65 °C, 15 min in a thermocycler for cell lysis. After a brief centrifugation, 1 μl of 10% tween-20 was added per well to quench SDS, then 4 μl of Q5 High-Fidelity 2X Master Mix (NEB, cat. no. M0541L) was added to each well. Pre-PCR was performed as follows: 72 °C 5 min, 98 °C 1 min; then 10 cycles of: 98 °C 20 s, 63 °C 20 s, 72 °C 1 min; and hold at 10 °C.

All pre-libraries in one 384-well plate were pooled into a 15 ml tube, and purified using Zymo DNA Clean & Concentrator kit (Zymo, cat. no. D4014). Excessive primers were digested by thermolabile EXO I, then the pre-libraries were purified using 1.2× VAHTS DNA clean beads (Vazyme, cat. no. N411-01) and elute in 15 μl Nuclease-free Water.

### GEMs generation and library pre-amplification in 10x droplet workflow

After EXO I digestion and gap fill-in, pre-libraries in the droplet workflow were generated using the 10x Genomics Chromium Controller Instrument (10x Genomics, Pleasanton, CA) and Chromium Next GEM Single Cell ATAC Library & Gel Bead Kit v1.1 following manufacturer recommended protocol, but with several modifications: 1) The transposition step using 10x ATAC enzyme is omitted (step 1); 2) After SPRIselect, DNA were eluted in 35.5 μl of Elution Solution I (step 3.2); 3) In sample index PCR, custom primers were used instead of SI-PCR primer B and individual Single Index N Set A (step 4.1); 4) After sample index, the 100 μl reaction was equally divided into two parts and purified separately for RNA and ATAC pre-libraries (step 4.2).

Briefly, nuclei were washed twice with 150 μl of ice-cold 1x DNB-0.5%BSA, resuspended in 15 μl of 1× DNB-0.5%BSA and counted. X μl of nuclei (3000~8000 cells) were mixed with 7 μl ATAC buffer B, and (8-X) μl of Qiagen buffer EB, to bring the total volume to 15 μl. Then, 60 μl of master mix was mixed with 15 μl transposed nuclei, and loaded on a Chromium controller Single-Cell Instrument to generate single-cell Gel Bead-In-Emulsions (GEMs). After breaking the GEMs, the barcoded DNA was purified using MyOne beads, size selected using 1.2x VAHTS DNA clean beads (Vazyme, cat. no. N411-01), and eluted in 35.5 μl of Elution Solution I. For sample index library pre-amplification, 35 μl of purified products was mixed with 50 μl Amp Mix, 5 μl Illumina P5 primer (10 μM), 5 μl RNA_droplet_N7xx primer (10 μM), and 5 μl ATAC_droplet_N7xx primer (10 μM). PCR was performed as follows: 98 °C 1 min; 7 cycles of: 98 °C 20 s, 63 °C 20 s, 72 °C 20 s; 1 cycle of 72 °C 1 min; then hold at 10 °C. 50 μl of the PCR product was taken for ATAC pre-library, purified with 1.0x VAHTS DNA clean beads (Vazyme), then eluted in 40.5 μl nuclease-free water. The remaining 50 μl for RNA pre-library was purified with 0.8x VAHTS DNA clean beads, and eluted in 40.5 μl Nuclease-free water.

### Final library amplification

For final ATAC+RNA library amplification in the plate-based workflow, 15 μl of purified pre-library was mixed with 2.5 μl RNA_plate_S5xx primer, 2.5 μl ATAC_plate_S5xx primer, 5 μl Illumina P7 primer, and 25 μl Q5 High-Fidelity 2× Master Mix. PCR was performed as follows: 98 °C 1 min; 10 cycles of: 98 °C 20 s, 63 °C 20 s, 72 °C 1 min; 1 cycle of 72 °C 5 min; then hold at 10 °C. Final library was purified using 1.2× VAHTS DNA cleaning beads and elute in 20 μl Nuclease-free Water.

For final ATAC library amplification in the 10x droplet workflow, 40 μl of purified 10x ATAC pre-library was mixed with 50 μl Q5 High-Fidelity 2× Master Mix, 5 μl Illumina P5 primer (10 μM), and 5 μl ATAC_droplet_N7xx primer (10 μM). PCR was performed as follows: 98°C 1 min; 7 cycles of: 98 °C 20 s, 63 °C 20 s, 72 °C 20 s; 1 cycle of 72 °C 1 min; then hold at 10 °C. Final ATAC library was purified using 1.0× VAHTS DNA cleaning beads and elute in 20 μl Nuclease-free Water.

For final RNA library amplification in the 10x droplet workflow, 40 μl of purified 10x RNA pre-libraries was mixed with 50 μl Q5 High-Fidelity 2X Master Mix, 5 μl Illumina P5 primer (10 μM), and 5 μl RNA_droplet_N7xx primer (10 μM). PCR was performed as follows: 98 °C 1 min; 7 cycles of: 98 °C 20 s, 63 °C 20 s, 72 °C 20 s; 1 cycle of 72 °C 1 min; then hold at 10 °C. Final RNA library was purified using 0.8× VAHTS DNA cleaning beads and elute in 20 μl Nuclease-free Water.

### 10x Multiome library preparation

K562, NIH3T3 and E14 cell nuclei were isolated according to 10x Genomics Demonstrated Protocol CG000365 Rev C. Libraries were generated on 10x Chromium Single-Cell Multiome ATAC + Gene Expression platform following the manufacturer’s protocol CG000338 Rev E.

### Sequencing

All ISSAAC-seq libraries were sequenced on Illumina NextSeq 500 and NovaSeq 6000 using standard protocols. For the plate-based libraries, 150 x 8 x 8 x 150 or 75 x 8 x 8 x 75 cycles was used. For the droplet-based libraries, 150 x 8 x 16 x 150 cycles was used. All 10x Multiome libraries were sequenced on Illumina NovaSeq 6000 following the recommendation from the 10x Genomics.

### Sequencing Data Pre-processing

FastQ files were generated using the bcl2fastq software (v2.20) from Illumina with the “-- create-fastq-for-index-reads” flag, which gave rise to four FastQ files per sample: Read 1, Read 2, Index 1 (i7) and Index 2 (i5). Samples were demultiplexed using deML^48^. For the plate-based workflow ATAC-seq data, Read 1 and Read 2 FastQ files were processed using the Snakemake^49^ pipeline as described previously^34^, except that BWA^50^ was used for the alignment. For the plate-based workflow RNA-seq data, the combination of i7 and i5 defines a single cell and the first 10 bp of Read 1 were UMIs. Therefore, sequences from Index 1, Index 2 and the first 10 bp of Read 1 were concatenated using a custom python script to generate a new FastQ file containing the cell barcode and UMI (CB_UMI.fastq). Then the CB_UMI.fastq and Read 2 files were used to make the gene expression matrix by STARsolo^51^. For the droplet-based workflow of ATAC-seq data, Read 1, Read 2 and Index 2 files were processed by Cell Ranger ATAC v2.0.0 from 10x Genomics. For the droplet-based workflow of RNA-seq data, the first 10 bp of Read 2 are UMIs. Therefore, sequences from Index 2 and the first 10 bp of Read 2 were concatenated to generate a new FastQ file containing the cell barcode and UMI (CB_UMI.fastsq). Then the CB_UMI.fastq and Read 1 files were used to generate the gene expression matrix by STARsolo^51^.

### Data quality comparisons to other methods

The code used to perform the comparison and generate the figures can be found in the GitHub repository (https://github.com/dbrg77/ISSAAC-seq). Briefly, ISSAAC-seq data were downsampled to 50,000 read pairs per cell (ATAC) or 50,000 reads per cell (RNA). For public data set, fastq files or count matrices were downloaded. The ATAC-seq reads were mapped using chromap^52^ and the RNA-seq reads were analysed using STARsolo^51^. The resulting count matrices were used to generate the quality metrics, including the number of reads in peaks, the number UMIs and the number of detected genes.

### Species mixing experiment

Briefly, 50,000 HEK293T and 50,000 NIH3T3 cells were combined and went through the ISSAAC-seq procedures. The resulting data were mapped to a GRCh38 and mm10 combined reference genome using chromap^52^ (ATAC) and STARsolo^51^ (RNA). Reads mapped to each genome in each cell barcode were counted. For ATAC data, we assign cell barcodes with >90% reads mapped to a single genome as singlets; for RNA data, we assign cell barcodes with >80% reads mapped to a single genome as singlets. For the joint analysis, we only treat cell barcodes as singlets where both modalities provide concordant results. Otherwise, they were labelled as doublets. The code can be found in the GitHub repository (https://github.com/dbrg77/ISSAAC-seq).

### Mouse cortex ISSAAC-seq RNA Data Analysis

For scRNA-seq analysis, the gene expression UMI matrix was normalized in the Seurat package^24^, using the “NormalizeData” function. Principal component analysis (PCA) was used to reduce the dimension of the data. The first 30 PCs were used to find the cell clusters using a modularity-based community detection algorithm^53^, and the final results were visualized using uniform manifold approximation and projection (UMAP)^37^. The marker genes from each cluster were identified using the “FindAllMarkers” function in the Seurat package that used the Wilcoxon rank-sum test to find significantly different genes. The average of expression of selected marker genes and the proportion of expressing cells within each cluster were plotted using the Scanpy package^54^. To annotate the cell types, scRNA-seq data from a previous study in the mouse cortex^38^ was used as a reference and passed to the “FindTransferAnchors” function in the Seurat package. Cell types for cells produced in this study were predicted using the “TransferData” function. Then marker genes from each cluster were taken into account to refine the automated prediction.

### Mouse cortex ISSAAC-seq ATAC Data Analysis

ATAC-seq data from the two biological replicates were merged together using the “aggr” function from the Cell Ranger ATAC software. The combined peak cell matrix was used for dimensionality reduction and clustering using the Signac package^55^. To find out the difference of the chromatin status between ATAC clusters A2, A3 and A9 within the Ex-L4/5 IT neurons (RNA cluster R3), reads from cells that belong to RNA cluster R2 and ATAC cluster A2, RNA cluster R2 and ATAC cluster A3, or RNA cluster R2 and ATAC cluster A9 were aggregated, and peak calling was performed on the aggregated reads using MACS2^56^. Then the top 1000 cluster specific peaks, sorted by fold enrichment from MACS2, were taken for motif analysis using HOMER^57^ and for the comparison with the histone modifications from the PairedTag data.

### Pseudotime Analysis

Cells from RNA cluster R20 (oligodendrocyte progenitor cells, OPC) and R21 (oligodendrocytes, Oligo) were taken out for pseudotime analysis. Differentially expressed genes between R20 and R21 (q value < 0.05) were used to construct the oligodendrocyte maturation trajectory using Monocle2^45^. The gene activity scores inferred from scATAC-seq data were produced using Signac. The dynamics of gene expression and chromatin accessibility of the differentially expressed genes were visualised side-by-side along the trajectory.

### Data availability

The raw sequencing data has been deposited at ArrayExpress under the accession number E-MTAB-11264.

### Code availability

The code used for the data processing and analysis mentioned in the method section is available on the GitHub repository https://github.com/dbrg77/ISSAAC-seq.

