## Supplementary Information for "ISSAAC-seq enables sensitive and flexible multimodal profiling of chromatin accessibility and gene expression in single cells"

Supplementary Tables 1 & 2

Supplementary Notes 1 – 4

Supplementary Reference

Supplementary Protocols 1 & 2

#### Supplementary Table 1

| Public data | Method | Modality | Public accessions | Total raw reads (fastq) | Number of provided cell barcodes | Average raw read per cell |
|---|---|---|---|---|---|---|
| K562 | SNARE-seq | ATAC | SRR8528319 | 57,171,705 | From the cell line mixture experiment. There are a total of 1,047 cells, of which 198 are K562. | 54,605 |
| | | RNA | SRR8528318 | 5,174,960 | From the cell line mixture experiment. There are a total of 1,631 cells, of which 355 are K562. | 3,172 |
| | SHARE-seq | ATAC | SRR10428394 | 16,168,027 | 5,220 | 3,097 |
| | | RNA | SRR10428405 | 4,338,379 | 7,755 | 559 |
| GM12878 | SNARE-seq | ATAC | SRR8528319 | 57,171,705 | From the cell line mixture experiment. There are a total of 1,047 cells, of which 138 are GM12878. | 54,605 |
| | | RNA | SRR8528318 | 5,174,960 | From the cell line mixture experiment. There are a total of 1,631 cells, of which 280 are GM12878. | 3,173 |
| | SHARE-seq | ATAC | SRR10428391 | 188,612,680 | 7,927 | 23,794 |
| | | RNA | SRR10428402 | 85,545,075 | 5,346 | 16,002 |
| | 10x Multiome | ATAC | SRR13716275 | 229,210,160 | 3,509 | 65,321 |
| | | RNA | SRR13716274 | 481,428,307 | 4,022 | 119,699 |
| H1 | SNARE-seq | ATAC | SRR8528319 | 57,171,705 | From the cell line mixture experiment. There are a total of 1,047 cells, of which 390 are H1. | 54,605 |
| | | RNA | SRR8528318 | 5,174,960 | From the cell line mixture experiment. There are a total of 1,631 cells, of which 471 are K562. | 3,173 |
| HEK293T | Paired-seq | ATAC | SRR8980188 | 164,634,027 | From the cell line mixture experiment. There are a total of 2,421 cells, of which 1,357 are HEK293T cells. | 68,002 |
| | | RNA | SRR8980189 | 159,005,510 | From the cell line mixture experiment. There are a total of 2,676 cells, of which 1,283 are HEK293T cells. | 59,419 |
| | sci-CAR-seq | ATAC | SRR7520209, SRR7520210, SRR7520211 | 429,236,798 | From the cell line mixture experiment. There are a total of 4,100 cells, of which 465 are HEK293T cells. | 104,692 |
| | | RNA | SRR7519729-SRR7520208 | 315,045,670 | From the cell line mixture experiment. There are a total of 5,581 cells, of which 465 are HEK293T cells. | 56,450 |
| NIH3T3 | SHARE-seq | ATAC | SRR10428393 | 16,713,613 | 2,159 | 7,741 |
| | | RNA | SRR10428404 | 9,090,473 | 2,314 | 3,928 |
| | sci-CAR-seq | ATAC | SRR7520209, SRR7520210, SRR7520211 | 429,236,798 | From the cell line mixture experiment. There are a total of 6,085 cells, of which 711 are NIH3T3 cells. | 70,540 |
| | | RNA | SRR7519729-SRR7520208 | 315,045,670 | From the cell line mixture experiment. There are a total of 6,093 cells, of which 711 are NIH3T3 cells. | 51,706 |

#### Supplementary Table 2

| Method | Sample | Modality | Public accessions | Total raw reads (fastq) | Number of provided cell barcodes | Average raw read per cell |
|---|---|---|---|---|---|---|
| SNARE-seq | Frozen adult cortex | ATAC | SRR9672090<br>SRR9672092<br>SRR9672094<br>SRR9672096<br>SRR9672098<br>SRR9672100<br>SRR9672102<br>SRR9672104<br>SRR9672106<br>SRR9672108<br>SRR9672110<br>SRR9672112 | 1,360,688,568 | 9,917 | 137,207 |
| | | RNA | SRR9672089<br>SRR9672091<br>SRR9672093<br>SRR9672095<br>SRR9672097<br>SRR9672099<br>SRR9672101<br>SRR9672103<br>SRR9672105<br>SRR9672107<br>SRR9672109<br>SRR9672111 | 1,933,856,377 | 24,074 | 80,329 |
| | Frozen P0 cortex | ATAC | SRR8528321<br>SRR8528323<br>SRR8528325<br>SRR8528327<br>SRR8528329 | 1,125,406,701 | 5,080 | 221,536 |
| | | RNA | SRR8528320<br>SRR8528322<br>SRR8528324<br>SRR8528326<br>SRR8528328 | 59,707,164 | 12,682 | 4,708 |
| Paired-seq | Frozen adult cortex | ATAC | SRR8980190<br>SRR8980192<br>SRR8980194 | 287,711,316 | 10,854 | 26,507 |
| | | RNA | SRR8980191<br>SRR8980193<br>SRR8980195 | 347,495,286 | 12,406 | 28,010 |
| | Frozen fetal brain | ATAC | SRR8980196<br>SRR8980198 | 226,642,404 | 18,554 | 12,215 |
| | | RNA | SRR8980197<br>SRR8980199 | 202,16,849 | 16,849 | 11,998 |
| SHARE-seq | Frozen mouse brain | ATAC | SRR10428398 | 38,283,279 | 2,475 | 15,467 |
| | | RNA | SRR10428409 | 98,103,936 | 3,107 | 31,575 |
| 10x Multiome | Fresh E18 mouse brain cortex | ATAC | https://www.10xgenomics.com/resources/datasets/fresh-embryonic-e-18-mouse-brain-5-k-1-standard-2-0-0 | 193,923,828 | 4,866 | 39,853 |
| | | RNA | | 474,002,632 | 6,049 | 78,360 |

**Supplementary Notes**

#### 1. Improvement of ISSAAC-seq

One of the key steps in ISSAAC-seq is to perform the chromatin tagging step at 30 °C, instead of the 37 °C that is commonly used in an ATAC-seq experiment. Chromatin tagging at 30 °C does not affect the complexity of the ATAC-seq library and increases the quality of the RNA-seq library significantly. Using the mouse E14 embryonic stem cell line, we see a 5- to 7-fold increase in the number of detected UMIs and 3 to 7 times more detected genes (**Supplementary Fig. 3a** and **b**). We also tested the effect of two commonly used RNase inhibitors: Protector RNase (Sigma, cat. no. 3335399001) recommended by 10x Genomics for the Multiome kit usage (https://kb.10xgenomics.com/hc/en-us/articles/360049543672-Can-I-use-an-alternative-RNase-inhibitor-part-number-) and RiboLock (Thermo, cat no. EO0382). The choice of the RNase inhibitor does have an effect, but the chromatin tagging temperature is the major factor for getting a high quality RNA-seq library.

During the *in situ* reverse transcription (RT) step, we included 12% polyethylene glycol (PEG) 8000 as a crowding reagent, which has been shown to increase the efficiency of the RT reaction in methods like mcSCRB-seq[1], SMART-seq3[2] and SHARE-seq[3]. In addition, the use of Tn5 transposase for the DNA/RNA hybrid tagging is efficient. This step also removes the necessity to perform a full-length cDNA synthesis and amplification using a template switch oligo (TSO), which can be tricky in certain situations, as shown in the method of Seq-Well S3[4]. Finally, ISSAAC-seq has fewer experimental steps compared to the previous "split-pool" based methods, which help to minimise material loss during the whole procedure.

In summary, we think the good performance of ISSAAC-seq may come from a combination of the aforementioned factors.

#### 2. Robustness of ISSAAC-seq

The ISSAAC-seq experiments are robust and reproducible. Cells from different experiments are well mixed on a UMAP space (**Supplementary Fig. 7** and **9**). They form clear clusters that can be identified as well-known cell types. The results in the paper are presented as they are, without applying any integration or batch correction methods. There seems to be some minor batch difference in the In-*Scn5a* (red dotted circle below) and Ex-PIR *Ndst4* clusters (blue dotted circle below):

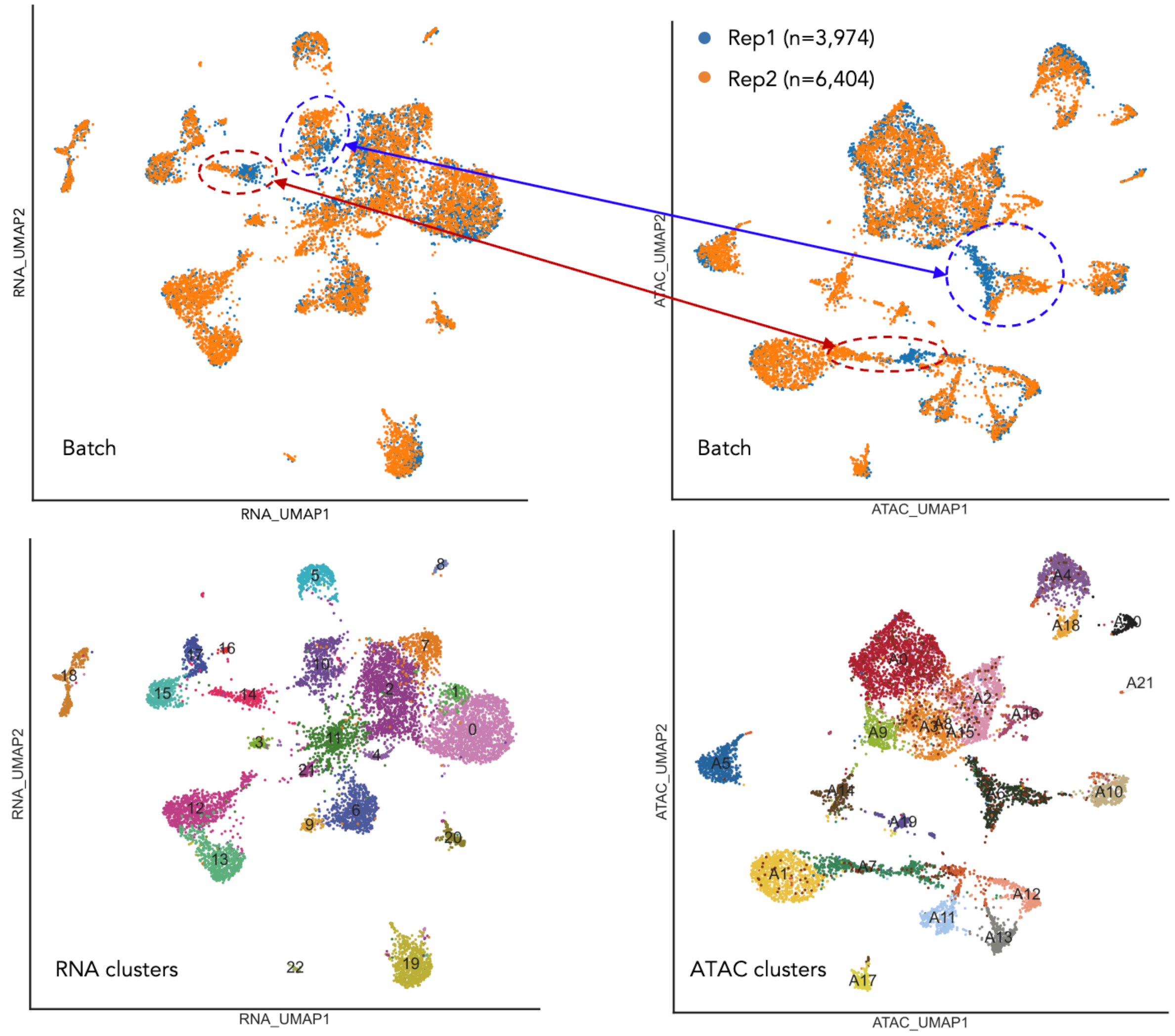


However, cells from different batches still stay together and the clustering results still suggest they are from the same clusters. Differential analysis on the Ex-PIR *Ndst4* neurons (blue dotted circles) returns no significant peaks and only 14 significant genes (q value < 0.01 and |log2FC| > 1) between the two batches. For the In-*Scn5a* (red dotted circles) cells, only 26 differentially expressed genes were found between the two batches, and they are mainly associated with functions involved in the regulation of neurotransmitter, synapse and metabolic processes *etc*.:

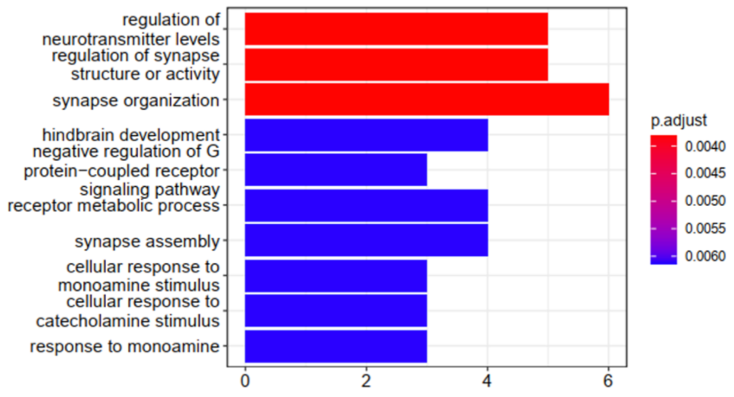

It is possible that those two cell types (In-*Scn5a* and Ex-PIR *Ndst4*) are prone to be affected by cell handling during the experiments. In addition, if we check the percentage of cells from each replicate per cluster, we found they roughly follow the expectation. The red dotted line in the figure below indicates 38% which is the percentage of replicate 1 cells in the whole data. Cells from both replicates are present in all clusters, with replicate 2 having roughly 1.6 times as many cells as replicate 1 in most clusters:

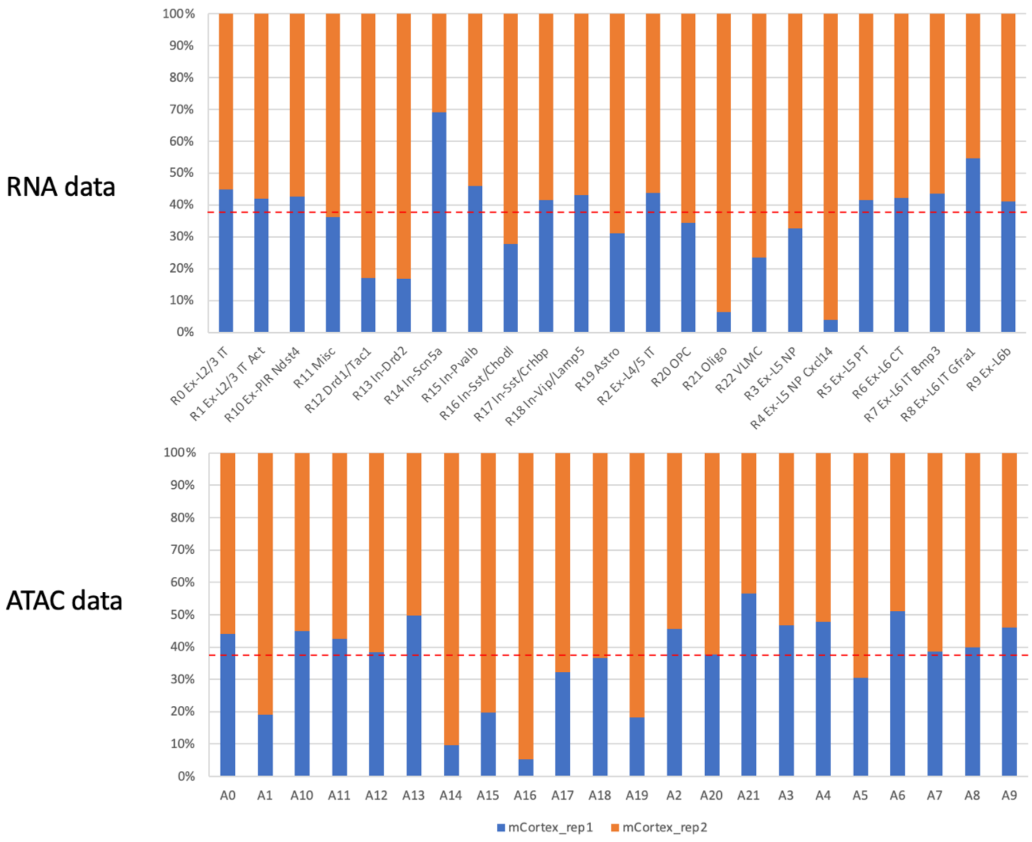


Finally, if an integration method was applied, such as Harmony[5], we can remove the separation of the batches from those two cell clusters:

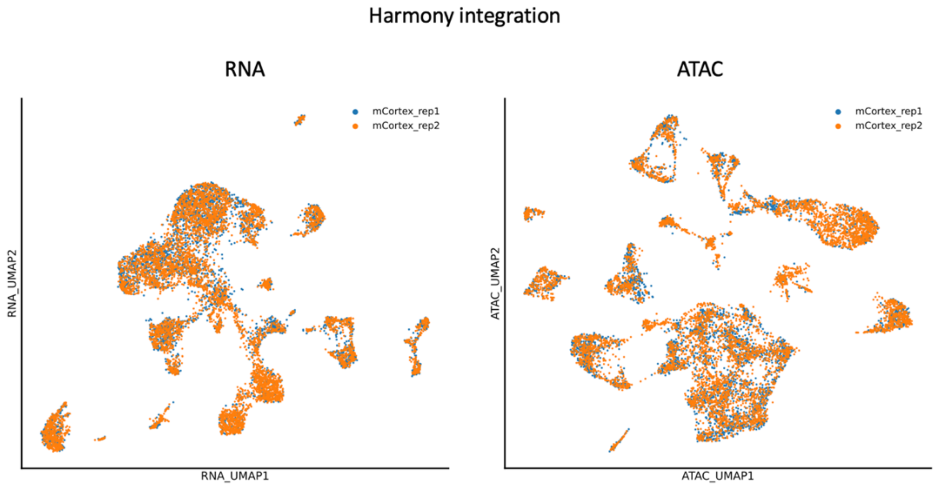

Since most cells from two different replicates are already well-mixed even before the integration, we did not use it in the paper to avoid over-integration. Based on the above results, ISSAAC-seq is robust and reproducible.

#### 3. Motivations of using ISSAAC-seq

**Cost benefits:** The 10x Genomics Single Cell Multiome ATAC + Gene Expression kit, hereafter referred to as the 10x Multiome kit, offers a commercial solution for the joint profiling of chromatin accessibility and gene expression from the same cell. Compared to the commercial solution, ISSAAC-seq achieves joint profiling of chromatin accessibility and gene expression from the same cell using only a small fraction of the cost of the 10x Mulitome kit, particularly for the plate-based workflow. The cost will increase if we adopt commercial kit for cell capture. In the most expensive case, 10x scATAC-seq kit was used for cell capturing and barcoding in ISSAAC-seq, and the total cost of ISSAAC-seq reaches about a half of the 10x Multiome kit. In addition, as an open-source method, all the reagents in ISSAAC-seq have been clearly listed. Users can further optimise the upfront reaction and potentially change to a different droplet-based system. Here, we only tested the 10x Geonomics platform. However, any systems with a Nextera Read 1 sequence on the bead should work, such as HyDrop[6] or Bio-Rad[7].

**Modular design:** ISSAAC-seq has a flexible and modular protocol design. Prior to single cell isolation, everything is done in nuclei using a population of cells, which allows easy handling of a large number of cells. All reagents in those steps can be further customised and modified in a sample specific manner if needed. In the current version, we have provided an initial condition that works well for the cell lines and primary tissues tested in the main text. The reaction condition of ISSAAC-seq is built upon the optimisation of single cell protocols developed in recent years, such as scTHS-seq[8], mcSCRB-seq[1], SPLiT-seq[9], SMART-seq3[2] and SHARE-seq[3].

**Easy and straightforward small-scale proof-of-principle tests:** At the stage of single cell isolation, the user can choose the method based on the project. If a large number of cells are not really needed, the plate-based workflow is a better option. This is useful to, for example, samples with limited number of available cells, small-scale projects and pilot testing for a new type of sample that the user is working on for the first time. This can also be combined with FACS index sorting to record extra information for each cell. The user can also use this workflow to test experimental conditions during the *in situ* reaction module without spending too much time and money. Typically, a 96- or 384-well plate is more than enough to tell if the experiment works or not. If a lot of cells are needed, the user can simply choose a droplet system with a Nextera Read 1 sequence.

#### 4. Further development of ISSAAC-seq

Recently, methods such as TEA-seq[10], DOGMA-seq[11] and NEAT-seq[12] take a step further to profile ATAC, RNA and protein information in the same cell by using antibody-oligo conjugates. This enables even more detailed characterization of cellular states. With the modular and flexible design of ISSAAC-seq, it should be straightforward to incorporate the same strategy to achieve similar outcomes. One possibility is to attach barcoded oligos with UMIs and partial Nextera Read 1 sequence to the antibody. In this case, it can be used directly in the current

ISSAAC-seq workflow, where the 10x Genomics Single Cell ATAC kit should be able to capture the antibody-oligo information due to the presence of the Nextera Read 1 sequence on the beads. However, attaching custom oligos to antibodies is difficult for many labs. Therefore, a more practical solution is to design a bridging oligo like that in DOGMA-seq[11] that brings the oligos on the commercial antibody to the bead from the 10x Genomics Single Cell ATAC kit. In this case, one side of the bridging oligo should be reverse complementary to the partial Nextera Read 1 sequence, and the other side should be reverse complementary to the oligo on the commercial antibodies, such as Total-SeqA/B by BioLegend.

On a related note, the antibody-oligo conjugates or lipid-modified oligos[13] can be used as hash tag oligos (HTOs) to merely label cells from different samples. Then cells can be "over-loaded" on the droplet machine, and multiplets can be removed by looking at HTOs. In this way, multiple samples and more cells can be done in one run to reduce the running cost. Furthermore, recent studies have incorporated the combinatorial indexing idea[14,15] with droplet technologies, such as dsciATAC-seq[7] and scifi-RNA-seq[16]. Similarly, we can use barcoded Tn5 for the chromatin and DNA/RNA hybryid tagging in ISSAAC-seq to label cells from dozens of samples. Then cells can be pooled and over-loaded on the 10x droplet machine, and single cells can be identified using a combination of barcodes on the 10x beads and the Tn5 barcodes. This should work in theory and will further increase the throughput and reduce the cost.

### Supplementary Protocol 1

#### ISSAAC-seq_10x_droplet_ workflow

Wei Xu, XiChen_lab@SUSTech, 2022.05

#### Reagents

- DNA-OFF (Takara Bio, cat. no. 9036)
- 1X DPBS (ThermoFisher, cat. no. 14190169)
- Trypan Blue Solution, 0.4% (ThermoFisher, cat. no.15250061)
- 30% (w/v) BSA (Sigma, cat.no. A8577)
- 5 M NaCl (Sigma, cat.no. S6546)
- 1 M $MgCl_2$ (ThermoFisher, cat. no. AM9530G)
- 1 M Tris-HCl, pH 8.0 (ThermoFisher, cat. no. 15568025)
- 1 M Tris-HCl, pH 7.5 (ThermoFisher, cat. no. 15567027)
- Glycerol (Sigma, cat.no. G5516)
- Triton X-100 (Sigma, cat.no. 93443-100ML)
- IGEPAL CA-630 (Sigma, cat. no. I8896)
- Tween-20 (Sigma, cat.no. 655205-250ML)
- 2% Digitonin (Promega, cat. no. G9441) **CRITICAL** Avoid more than 5 freeze thaw cycles.
- N,N-Dimethylformamide (Sigma, cat. no. D4551-250ML)
- PEG8000 (Sigma, cat.no. 89510-250G-F)
- 1 M DTT (Sigma, cat.no. 43816-10ML)
- 5 M Potassium acetate (Sigma, cat. no. 95843-100ML-F)
- 1 M Magnesium acetate (Sigma, cat. no. 63052-100ML)
- 0.5 M EDTA, pH 8.0 (ThermoFisher, cat. no. AM9260G)
- dNTP Set 100 mM Solutions (ThermoFisher, cat.no. R0181)
- RiboLock RNase Inhibitor (ThermoFisher, cat.no. EO0382)
- RNaseOUT™ Recombinant Ribonuclease Inhibitor (ThermoFisher, cat.no. 10777019)
- SUPERase•In™ Rnase Inhibitor (ThermoFisher, cat.no. AM2694)
- Maxima H Minus Reverse Transcriptase (ThermoFisher, cat.no. EP0753)
- Thermolabile Exonuclease 1 (NEB, cat.no. M0568)

- Ethanol (Sangon Biotech, cat. no. A500737-0500)
- Chromium Next GEM Single Cell ATAC Library & Gel Bead Kit v1.1 (10x Genomics, cat.no.1000175)
- Chromium Next GEM Chip H Single Cell Kit (10x Genomics, cat.no.1000161)
- Q5 High-Fidelity 2X Master Mix (NEB, cat.no. M0492L)
- VAHTS® DNA Clean beads (Vazyme, cat. no. N411-01)
- UltraPure™ DNase/RNase-Free Distilled Water (Invitrogen, cat. no. 10977015)
- Qubit™ dsDNA HS Assay Kit (ThermoFisher, cat.no. Q32851)
- High Sensitivity DNA Kit (Agilent, cat. no. 5067-4626)
- Primer oligos. All oligos were ordered from Sangon Biotech (Shanghai) with standard purification.

#### Equipment

- 1.5 mL Eppendorf Safe-Lock Tubes (Eppendorf, cat. no. 0030120086)
- 0.2 mL PCR Tubes (Gunster Biotech, cat. no. MB-P08-A)
- Qubit™ Assay Tubes (ThermoFisher, cat.no. Q32856)
- C-Chip disposable hemocytometer (Incyto, cat. no. DHC-N01-5)
- Centrifuge (Eppendorf, centrifuge 5424R)
- ThermoMixer C (Eppendorf, cat. no. 5382000074)
- Alpha Thermal Cycler, with 96-well and 384-well blocks (PCRmax)
- Chromium Controller & Next GEM Accessory Kit (10x Genomics, cat. no. 1000204)
- 10x Vortex Adapter (10x Genomics, cat. no. 120251)
- Chromium Next GEM Secondary Holder (10x Genomics, cat. no. 1000195)
- 10x Magnetic Separator (10x Genomics, cat. no. 120250)
- Qubit 4 (ThermoFisher, cat.no. Q33238)
- 2100 Bioanalyzer Laptop Bundle (Agilent, cat. no. G2943CA)

#### Reagent setup

**Annealing buffer**

Mix 10 µl 1 M Tris-HCl (pH 8.0), 10 µl 5 M NaCl, 2 µl 0.5 M EDTA (pH=8.0), 978 µl Nuclease-free Water for 1 ml Annealing buffer. Stored at -20°C for up to 1 year.

**Coupling buffer**

Mix 50 µl 1 M Tris-HCl (pH 7.5), 20 µl 5 M NaCl, 2 µl 50 mM EDTA (pH=8.0), 10 µl 10% TritonX-100, 1 µl 1 M DTT, 500 µl glycerol, and 417 µl Nuclease-free Water for 1 ml coupling buffer. Stored at -20°C for up to 1 year.

**1× DPBS-0.5%BSA-RI**

Mix 166.7 µl 30% (w/v) BSA, 50 µl RiboLock RNase Inhibitor (40 U/µl), and 10 ml 1× DPBS. The buffer can be stored at -20°C for up to 2 weeks.

**0.5×DPBS-0.5%BSA-RI**

Mix 166.7 µl 30% (w/v) BSA,50 µl RiboLock RNase Inhibitor (40 U/µl), 5 ml 1× DPBS, and 5 ml Nuclease-free Water. The buffer can be stored at -20°C for up to 2 weeks.

**Omni-ATAC RSB**

Mix 500 µl 1 M Tris-HCl (pH 7.5), 100 µl 5 M NaCl, 150 µl 1 M $MgCl_2$, and 49.25 ml Nuclease-free Water for 50 ml omni-ATAC RSB. Sterilize through 0.22 µm filter, stored at 4°C for up to 2 months.

**RSB-DTN**

Mix 487.5 µl Omni-ATAC RSB, 5 µl 10% Tween-20, 5 µl 10% IGEPAL CA-630 and 2.5 µl 2% Digitonin. Make fresh.

**RSB-T**

Mix 10 ml Omni-ATAC RSB and 100 µl 10% Tween-20. Store at 4°C for up to 1 month.

**0.1% digitonin**

Mix 0.5 µl 2% digitonin stock with 9.5 µl Nuclease-free Water. **CRITICAL** Make fresh and avoid freeze thaw cycles. The Promega 2% digitonin stock is dissolved in DMSO. Thaw the digitonin stock at room temperature.

**4× THS TD buffer**

Mix 132 µl 1 M Tris-HCl (pH8.0), 52.8 µl 5 M Potassium acetate, 40 µl 1 M Magnesium acetate, 640 µl N, N-Dimethylformamide (DMF) and 135.2 µl Nuclease-free Water for 1 ml 4× THS TD buffer. Store at -20°C for up to 2 months.

**2× Stop Buffer**

Mix 500 µl 1x PBS, 40 µl 0.5M EDTA (pH8.0), 66 µl 30% BSA and 394 µl Nuclease-free Water for 1 ml 2× Stop Buffer. Make fresh.

**50% (w/w) PEG8000**

Mix 5 g PEG8000 and 5 ml Nuclease-free Water, sterilize through 0.22 µm filter, aliquot and store at -20°C for up to 2 months.

**5× NaCl RT Buffer**

Mix 1.25 ml Tris-HCl (pH =8.0), 375 µl 5 M NaCl, 75 µl MgCl2, 250 µl 1 M DTT, 3.05 ml Nuclease-free Water to prepare 5 ml 5× NaCl RT Buffer. Sterilize through 0.22 µm filter, aliquot and store at -20°C for up to 2 months.

**80% (v/v) ethanol**

Mix 4 ml absolute ethanol with 1 ml Nuclease-free Water. Make fresh.

**1x DNB-0.5%BSA**

Mix 50 µl 20x DNB (10x genomics), 16.7 µl 30% BSA and 933.3 µl Nuclease-free Water. Make fresh.

**50% (v/v) glycerol**

Mix 5 ml glycerol and 5 ml Nuclease-free Water. Sterilize through 0.22 µm filter, aliquot and store at 4°C for up to 3 months.

#### Procedures

**Prepare Tn5 transposome complexTiming 2.5 h**

1. Dissolve ME_S5, ME_S7 and ME_bottom oligos respectively in annealing buffer to a final concentration of 100 µM.
2. Mix 25 µl ME_S5 oligo (100 µM) with 25 µl ME_Bottom oligo (100 µM) in a PCR tube, and anneal in a thermocycler as follows: 98 °C for 3 min, and slowly cool down to 16 °C with a temperature ramp of −0.1 °C/s, to generate S5_adaptor (50 µM).
3. Similarly, mix 25 µl ME_S7 oligo (100 µM) with 25 µl ME_Bottom oligo (100 µM) in a PCR tube and anneal to form S7_adaptor (50 µM).
4. Dilute the annealed 50 µM S5_adaptor or S7_adaptor with Nuclease-free Water to 20 µM.
5. To assemble Tn5-S5/S7 transposome, add 12 µl S5_adaptor (20 µM), 12 µl S7_adaptor (20 µM), 48 µl purified Tn5 (0.5 µg/µl), and 88 µl coupling buffer to a 1.5 ml tube, mix thoroughly by gently pipetting up and down 10 times. Incubate at room temperature for 1 h, then store at -20 °C.
6. To assemble Tn5-S5/S5 transposome, add 24 µl S5_adaptor (20 µM), 48 µl purified Tn5 (0.5 µg/µl), and 88 µl coupling buffer to a 1.5 ml tube, mix thoroughly by gently pipetting up and down 10 times. Incubate at room temperature for 1 h, then store at -20 °C.

PAUSE POINT The assembled Tn5 transposome could be stored at -20°C for up to 6 months.

**Prepare nuclei from cell lines Timing 1 h**

CRITICAL Work quickly and always keep the samples on ice unless otherwise indicated.

CRITICAL All centrifugation should be performed at 4 °C in a swing bucket centrifuge. DO NOT use fixed-angle centrifuges, which will cause severe cell loss during washes.

CRITICAL When removing the supernatant, be careful not to disturb the cell pellet.

7. Coat the tube: add 0.5 ml 1× DPBS–0.5%BSA-RI (0.2 U/µl) to a 1.5 mL tube, invert the tube three to five times, then aspirate all liquid in the tube and inside the cap and discard.

CRITICAL Cells may attach to the inner surface of the centrifugation tubes. Coating the tubes with 0.5% (w/v) BSA helps reduce cell loss when working with low amounts of cells.

8. Count cells using a C-Chip disposable hemocytometer and use Trypan Blue to check cell viability. Make sure more than 90% cells are viable.
9. Spin down 100,000 cells per sample, centrifuge at 500g, 4°C for 5 min.

**CRITICAL** Perform all centrifugations on a swing bucket centrifuge to reduce cell loss.

10. Wash the pellet once with 500 µl ice-cold DPBS-0.5% BSA-RI, centrifuge at 500g, 4°C, 5 min.
11. Resuspend cells in 50 µl ice-cold RSB-DTN- RI (0.8 U/µl) and leave on ice for 3 min.
12. Add 1 ml ice-cold RSB-T to the tubes (1.05 ml in total), invert 3 times to mix.
13. Centrifuge at 1000g, 4 °C for 8 min to collect the nuclei.
14. Aspirate all supernatants and hold on ice.

#### Chromatin tagmentation Timing 1 h

15. Prepare 50 µl chromatin tagmentation reaction for each sample：

| | Stock | Final | Volume (50 µl) |
|---|---|---|---|
| 4×THS TD buffer | 4× | 1× | 12.5 µl |
| 0.1% Digitonin | 0.1% | 0.01% | 5 µl |
| Tn5-S5/S7 | - | - | 2.5 µl |
| RiboLock RNase inhibitor | 40 U/µl | 1.2 U/µl | 1.5 µl |
| SUPERaseIn RNase inhibitor | 20 U/µl | 0.4 U/µl | 1 µl |
| RnaseOUT RNase inhibitor | 40 U/µl | 0.8 U/µl | 1 µl |
| Nuclease-free Water | - | - | 26.5 µl |

**CRITICAL** Tn5 should be the last component to add, *i.e.,* add it right before use.

16. Resuspend each pellet in 50 µl tagmentation mixture by gently pipetting up and down 30 times.

    **CRITICAL** Perform gentle and slow pipetting to avoid disrupting the nucleus structure. Make sure that the nuclei are intact after pipetting. Mix gently and avoid bubbles.
17. Incubate at 30°C, 800 rpm for 30 min.
18. Add 50 µl 2× Stop buffer to each sample and gently pipette up and down 6~8 times to mix.
19. Centrifuge at 1000g, 4 °C for 5 min. Remove the supernatants and wash the pellet twice with 200 µl ice-cold 0.5×PBS-0.5%BSA-RI.
20. Carefully remove all supernatants and hold on ice.

    **CRITICAL** The nuclei pellet may be invisible, be careful not to disrupt the nuclei at the bottom of the tube when aspirating the supernatant. It is okay to have 2–3 µl buffer leftover at this step.

**Reverse transcription Timing 1.5 h**

21. Prepare 100 µl RT reaction for each sample, slowly pipette up and down 10~20 times to mix.

| Reagent | Stock | Final | Volume (100 µl) |
|---|---|---|---|
| TruseqR2_oligo_dT | 10 µM | 2 µM | 20 µl |
| dNTP | 10 mM each | 0.5 mM each | 5 µl |
| Maxima H minus Reverse Transcriptase | 200 U/µl | 10 U/µl | 5 µl |
| RiboLock RNase Inhibitor | 40 U/µl | 2 U/µl | 5 µl |
| SUPERaseIn RNase Inhibitor | 20 U/µl | 0.2 U/µl | 1 µl |
| RnaseOUT RNase Inhibitor | 40 U/µl | 0.4 U/µl | 1 µl |
| 5× NaCl RT Buffer | 5× | 1× | 20 µl |
| 50% PEG8000 | 50% | 12% | 24 µl |
| Nuclease-free Water | - | - | 19 µl |

CRITICAL The 50% PEG8000 solution is viscous. Pipette slowly to ensure that no liquid remains along the tube sidewalls and tips.

22. Resuspend each pellet with 100 µl RT buffer by gently pipetting up and down 50 times.

CRITICAL Perform gentle and slow pipetting to avoid disrupting the nucleus structure. Make sure that the nuclei are intact after pipetting. Mix gently and avoid bubbles.

23. Transfer the 100 µl RT mixture to a new PCR tube and incubated in a thermocycler for 10 min at 50˚C before cycling for three times at 8˚C for 12s, 15˚C for 45s, 20˚C for 45s, 30˚C for 30s, 42˚C for 2 min, and 50˚C for 3 min, followed by a final step at 50˚C for 5 min.

24. Transfer the RT mixture to a new, BSA coated and chilled 1.5 ml tube.

25. Centrifuge at 1000g, 4 °C for 5 min. Remove the supernatants and wash the pellet twice with 200 µl ice-cold 0.5×DPBS-0.5%BSA.

26. Carefully remove all supernatants and hold on ice.

CRITICAL After reverse transcription, the nuclei will be partially lysed and the number of intact nuclei will decrease. The pellet may be invisible, be careful not to disrupt the nuclei at the bottom of the tube when aspirating the supernatant. It is okay to have 2–3 µl buffer leftover at this step.

**mRNA-cDNA hybrid tagmentation Timing 1 h**

27. Set up RNA-DNA hybrid tagmentation mix, prepare 50 µl for each sample:

| | Stock | Final | Volume (50 µl) |
|---|---|---|---|
| 4×THS TD buffer | 4× | 1× | 12.5 µl |
| 0.1% Digitonin | 0.1% | 0.01% | 5 µl |
| Tn5-S5/S5 | - | - | 1.5 µl |
| Nuclease-free Water | - | - | 31 µl |

28. Resuspend each pellet with 50 µl tagmentation buffer by gently pipetting up and down 30 times.

CRITICAL Perform gentle and slow pipetting to avoid disrupting the nucleus structure. Make sure that the nuclei are intact after pipetting. Mix gently and avoid bubbles.

29. Incubate at 37°C, 800rpm, 30 min.

30. During incubation, take out the 20X Nuclei Buffer, ATAC Buffer B, Reducing Agent B, and Barcoding Reagent B from -20 °C, and Single Cell ATAC Gel Beads v1.1 from -80 °C (10X genomics, Chromium Next GEM Single Cell ATAC Library Kit and Gel Bead Kit v1.1), equilibrate to room temperature.

CRITICAL Equilibrate to room temperature at least 30 min before loading the chip.

31. Add 50 µl 2× Stop Buffer to each sample, mix thoroughly by gentle pipetting 6~8 times.

32. Centrifuge at 1000g, 4 °C for 5 min. Remove the supernatants and wash the pellet twice with 200 µl ice-cold 0.5×DPBS-0.5%BSA.

33. Carefully remove all supernatants and hold on ice.

CRITICAL The pellet may be invisible, be careful not to disrupt the nuclei at the bottom of the tube when aspirating the supernatant. It is okay to have 2–3 µl buffer leftover at this step.

#### EXO I digestion and Gap fill-in Timing 30 min

34. Prepare 50 µl reaction for each sample:

| Reagent | Stock | Final | Volume (50 µl) |
|---|---|---|---|
| dNTP | 10 mM each | 0.5 mM each | 2.5 µl |
| Maxima H Minus Reverse Transcriptase | 200 U/µl | 8 U/µl | 2 µl |
| Exo1 | 20 U/µl | 2 U/µl | 5 µl |
| 5× NaCl RT Buffer | 5× | 1× | 10 µl |
| Nuclease-free Water | - | - | 30.5 µl |

35. Resuspend each pellet in 50 µl reaction by gently pipetting up and down 30 times.
36. Incubate at 37°C for 15 min.
37. Centrifuge at 1000g, 4 °C for 5 min, aspirate all supernatants.
38. Wash the pellet twice with 150 µl of ice-cold 1x DNB (10x genomics)-0.5%BSA. Centrifuged at 1000g 4 °C for 5 min.
39. Remove all supernatants and hold on ice.

#### GEM Generation and Barcoding Timing 2 h

**CRITICAL** The following steps are modified form "Chromium Next GEM Single Cell ATAC Reagent Kits v1.1 User Guide" (CG000209, 10x genomics).

40. Resuspend cells in ~15 µl of 1× DNB-0.5%BSA. Mix 2 µl cell suspension with 8 µl 0.4% trypan blue and count the nuclei using a C-Chip disposable hemocytometer.

    **CRITICAL** Set a proper resuspension volume based on the total cell number and targeted loading concentration.
41. Prepare "Transposed Nuclei": Add 7 µl ATAC buffer B, (8-x) µl Qiagen buffer EB, and x µl nuclei (5000 ~ 10000 nuclei) in order, to a chilled 1.5 ml tube. The total volume is 15 µl. Pipette mix 10 times using a 10 µl tip. Centrifuge briefly and hold on ice.
42. Prepare Master Mix: Add 56.5 µl Barcoding Reagent B, 1.5 µl Reducing Agent B, and 2 µl

Barcoding Enzyme in order, to a chilled 1.5 ml tube. Pipette mix ~ 10 times using a 10 µl tip. Hold on ice.

43. Assemble CHIP H following the 10x instructions.
44. Dispense 50% Glycerol into Unused Chip Wells.
45. Add 60 µl master mix to 15 µl transposed nuclei, Pipette mix 20 times. Using the same pipette tip, dispense 70 µl Master Mix + Transposed Nuclei into the bottom center of each well in row labeled 1 without introducing bubbles.
46. Load Gel beads and Partitioning oil to each well following the 10x instructions.
47. Attach 10x Gasket, run the chip in the controller immediately.
48. Transfer the 100 µl GEMs to a chilled PCR 8-tube strip following the 10x instructions.
49. GEM incubation: incubate the PCR strip in a thermal cycler with the following program.

| Temperature | Time | Number of cycles |
|---|---|---|
| 72°C | 5 min | 1 |
| 98°C | 30 sec | 1 |
| 98°C | 10 sec | 12 |
| 59°C | 30 sec | |
| 72°C | 1 min | |
| 15°C | ∞ | 1 |

**PAUSE POINT** The GEMs could be stored at 15°C for up to 18 h or at -20°C for up to a week.

#### Post GEM incubation Cleanup Timing 1 h

50. Follow "Step 3" in the Chromium Next GEM Single Cell ATAC Reagent Kits v1.1 User Guide (CG000209), but elute the final products in 35.5 µl Elution solution 1 (step 3.2 j).

**PAUSE POINT** The eluted products could be stored at 4°C for up to 72 h or at -20°C for up to 2 weeks.

#### Library construction Timing 2 h

51. Set up pre-amplification reaction as follows:

| | |
|---|---|
| 5 µl | ATAC_droplet_N7xx (10 µM) |
| 5 µl | Illumina P5 (10 µM) |
| 5 µl | RNA_droplet_N7xx (10 µM) |

| | |
|---|---|
| 50 µl | 10x Amp Mix |
| 35 µl | Purified DNA from above |

52. Incubate in a thermal cycler with the following protocol for pre-amplification:

| | Temperature | Time | Number of cycles |
|---|---|---|---|
| Initial denaturation | 98°C | 1 min | 1 |
| Denaturation | 98°C | 20 sec | 7 |
| Annealing | 63°C | 20 sec | |
| Extension | 72°C | 20 sec | |
| Final extension | 72°C | 1 min | 1 |
| Hold | 10°C | ∞ | 1 |

53. Final ATAC library amplification: Take 50 µl of the pre-PCR product as ATAC pre-library, and purify with 1.0x VAHTS DNA clean bead, then elute in 40.5 µl Nuclease-free Water.

**PAUSE POINT** The ATAC pre-library could be stored at 4°C for up to 72 h or at -20°C for up to 2 weeks.

54. Setup ATAC library amplification reaction as follows:

| | |
|---|---|
| 5 µl | ATAC_droplet_N7xx (10 µM) |
| 5 µl | Illumina P5 (10 µM) |
| 50 µl | Q5 High-Fidelity 2× Master Mix |
| 40 µl | Purified DNA from above |

**CRITICAL** The ATAC_droplet_N7xx primers have the ATAC library sample barcodes. Use the same ATAC_droplet_N7xx primer for the same sample in pre-amplification and final ATAC library amplification. Use different primers for different samples if you want to sequence the samples in the same lane.

55. Perform ATAC library PCR as follows:

| | Temperature | Time | Number of cycles |
|---|---|---|---|
| Initial denaturation | 98°C | 1 min | 1 |
| Denaturation | 98°C | 20 sec | 7 |
| Annealing | 63°C | 20 sec | |

| | | | |
|---|---|---|---|
| Extension | 72°C | 20 sec | |
| Final extension | 72°C | 1 min | 1 |
| Hold | 10°C | ∞ | 1 |

56. Purify the final ATAC library using 1.0× VAHTS DNA cleaning beads and elute in 20 µl Nuclease-free Water.
57. Final RNA library amplification: Purify the remaining 50 µl pre-PCR products from step 52) with 0.8x VAHTS DNA clean beads, and elute in 40.5 µl Nuclease-free Water.

    **PAUSE POINT** The RNA pre-library could be stored at 4°C for up to 72 h or at -20°C for up to 2 weeks.
58. Set up RNA library amplification reaction as follows:

| | |
|---|---|
| 5 µl | RNA_droplet_N7xx (10 µM) |
| 5 µl | Illumina P5 (10 µM) |
| 50 µl | Q5 High-Fidelity 2× Master Mix |
| 40 µl | Purified DNA from above |

**CRITICAL STEP** The RNA_droplet_N7xx primers have the RNA library sample barcodes. Use the same RNA_droplet_N7xx primer for the same sample in pre-amplification and final RNA library amplification. Use different primers for different samples if you want to sequence the samples in the same lane.

59. Performed RNA library PCR as follows:

| | Temperature | Time | Number of cycles |
|---|---|---|---|
| Initial denaturation | 98°C | 1 min | 1 |
| Denaturation | 98°C | 20 sec | |
| Annealing | 63°C | 20 sec | 7 |
| Extension | 72°C | 20 sec | |
| Final extension | 72°C | 1 min | 1 |
| Hold | 10°C | ∞ | 1 |

60. Purify the final RNA library using 0.8× VAHTS DNA cleaning beads and elute in 20 µl Nuclease-free Water.

**Library QC and sequencing Timing 4~6 d**

61. Measure the concentration of each library by Qubit according to the manufacturer's

instructions.

62. Dilute the libraries to 5~10 ng/μl using Nuclease-free Water.
63. Check the library size distribution on an Agilent high-sensitivity chip according to the manufacturer's instructions. A typical ISSAAC RNA library shows a broad peak between 250-1000 bp. A typical ISSAAC ATAC library shows a nucleosome ladder between 200-1000 bp.
64. Sequence the libraries for 150 x 8 x 16 x 150 cycles on Novaseq 6000 platform (Illumina).

#### Timing

| | |
|---|---|
| Prepare Tn5 transposome complex | 2 h 30 min |
| Prepare nuclei from cell lines | 1 h |
| Chromatin tagmentation | 1 h |
| Reverse transcription | 1 h 30 min |
| mRNA-cDNA hybrid tagmentation | 1 h |
| EXO I digestion and Gap fill-in | 30 min |
| GEM Generation and Barcoding | 2 h |
| Post GEM incubation Cleanup | 1 h |
| Library construction | 2 h |
| Library QC and sequencing | 4~6 d |

#### Anticipated Results

**<u>Step 61):</u>** The library concentration is usually around 5~30 ng/μl (depending on cell types), which is sufficient for sequencing.

**<u>Step 63):</u>** A typical ISSAAC RNA library shows a broad peak between 250-1000 bp, and typical ISSAAC ATAC library shows a nucleosome ladder between 200-1000 bp.

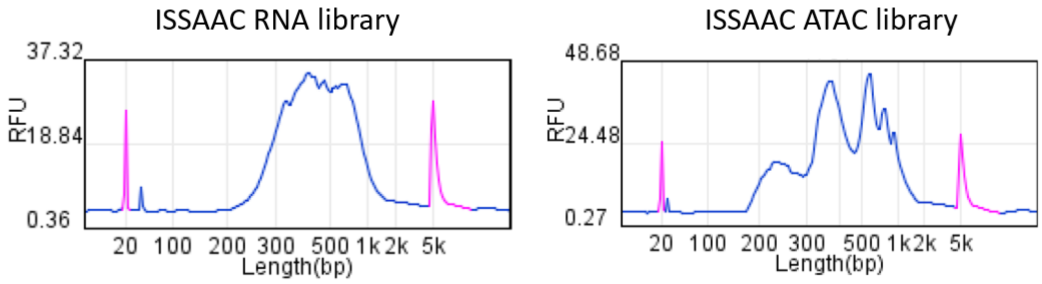

### Supplementary Protocol 2

#### ISSAAC-seq_FACS_workflow

Wei Xu, XiChen_lab@SUSTech, 2022.05

#### Reagents

- DNA-OFF (Takara Bio, cat. no. 9036)
- 1X DPBS (ThermoFisher, cat. no. 14190169)
- Trypan Blue Solution, 0.4% (ThermoFisher, cat. no.15250061)
- 30% (w/v) BSA (Sigma, cat.no. A8577)
- 5 M NaCl (Sigma, cat.no. S6546)
- 1 M $MgCl_2$ (ThermoFisher, cat. no. AM9530G)
- 1 M Tris-HCl, pH 8.0 (ThermoFisher, cat. no. 15568025)
- 1 M Tris-HCl, pH 7.5 (ThermoFisher, cat. no. 15567027)
- Glycerol (Sigma, cat.no. G5516)
- Triton X-100 (Sigma, cat.no. 93443-100ML)
- IGEPAL CA-630 (Sigma, cat. no. I8896)
- Tween-20 (Sigma, cat.no. 655205-250ML)
- 2% Digitonin (Promega, cat. no. G9441) **CRITICAL** Avoid more than 5 freeze thaw cycles.
- N,N-Dimethylformamide (Sigma, cat. no. D4551-250ML)
- PEG8000 (Sigma, cat.no. 89510-250G-F)
- 1 M DTT (Sigma, cat.no. 43816-10ML)
- 5 M Potassium acetate (Sigma, cat. no. 95843-100ML-F)
- 1 M Magnesium acetate (Sigma, cat. no. 63052-100ML)
- 0.5 M EDTA, pH 8.0 (ThermoFisher, cat. no. AM9260G)
- dNTP set 100 mM Solutions (ThermoFisher, cat.no. R0181)
- RiboLock RNase Inhibitor (ThermoFisher, cat.no. EO0382)
- RNaseOUT™ Recombinant Ribonuclease Inhibitor (ThermoFisher, cat.no. 10777019)
- SUPERase•In™ Rnase Inhibitor (ThermoFisher, cat.no. AM2694)
- Maxima H Minus Reverse Transcriptase (ThermoFisher, cat.no. EP0753)
- Thermolabile Exonuclease 1 (NEB, cat.no. M0568)

- DAPI (ThermoFisher, cat. no. 62248)
- 10% SDS (sigma, cat. no. L4509)
- DNA clean & concentrator-5 (Zymo, cat.no. D4014)
- Q5 High-Fidelity 2X Master Mix (NEB, cat.no. M0492L)
- VAHTS® DNA Clean beads (Vazyme, cat. no. N411-01)
- Ethanol (Sangon Biotech, cat. no. A500737-0500)
- UltraPure™ DNase/RNase-Free Distilled Water (Invitrogen, cat. no. 10977015)
- Qubit™ dsDNA HS Assay Kit (ThermoFisher, cat.no. Q32851)
- High Sensitivity DNA Kit (Agilent, cat. no. 5067-4626)
- Primer oligos. All oligos were orded from Sangon Biotech (Shanghai) with standard purification.

#### Equipment

- 1.5 mL Eppendorf Safe-Lock Tubes (Eppendorf, cat. no. 0030120086)
- 0.2 mL PCR Tubes (Gunster Biotech, cat. no. MB-P08-A)
- C-Chip disposable hemocytometer (Incyto, DHC-N01-5)
- Hard-Shell 384-Well PCR Plates (Bio-Rad, cat. No. HSP3801)
- Falcon 5 mL Round Bottom Polystyrene Test Tubes (for FACS sorting) (Corning, cat. no. 352054)
- Centrifuge (Eppendorf, centrifuge 5424R)
- Centrifuge for multi-well plates (Ke Cheng Equipment, L3-6K)
- 25 ml Reagent Reservoir (Biologix, cat. no. 25-1025)
- Low Dead Volume Reservoir/Collection Plate (for pooling libraries from 384-well plate) (Clickbio, cat. no. VBLOK200)
- PCR Sealing Film (Genstar, cat. no. HC96552-05)
- E1-ClipTip Bluetooth Electronic Multichannel Pipettes, 0.5 to 12.5µl (Thermo Scientific)
- 12.5µl ClipTip 384-Format Pipette Tips (Thermo Scientific, cat. no. 94410050)
- ThermoMixer C (Eppendorf, cat. no. 5382000074)
- Alpha Thermal Cycler, with 96-well and 384-well blocks (PCRmax)
- BD FACSAria sorter (BD Biosciences)
- Qubit 4 (ThermoFisher, cat.no. Q33238)

- 2100 Bioanalyzer Laptop Bundle (Agilent, cat. no. G2943CA)

#### Reagent setup

**Annealing buffer**

Mix 10 µl 1 M Tris-HCl (pH 8.0), 10 µl 5 M NaCl, 2 µl 0.5 M EDTA (pH=8.0), 978 µl Nuclease-free Water for 1 ml Annealing buffer. Stored at -20°C for up to 1 year.

**Coupling buffer**

Mix 50 µl 1 M Tris-HCl (pH 7.5), 20 µl 5 M NaCl, 2 µl 50 mM EDTA (pH=8.0), 10 µl 10% TritonX-100, 1 µl 1 M DTT, 500 µl glycerol, and 417 µl Nuclease-free Water for 1 ml coupling buffer. Stored at -20°C for up to 1 year.

**1× DPBS-0.5%BSA-RI**

Mix 166.7 µl 30% (w/v) BSA, 50 µl RiboLock RNase Inhibitor (40 U/µl), and 10 ml 1× DPBS. The buffer can be stored at -20°C for up to 2 weeks.

**0.5×DPBS-0.5%BSA-RI**

Mix 166.7 µl 30% (w/v) BSA, 50 µl RiboLock RNase Inhibitor (40 U/µl), 5 ml 1× DPBS, and 5 ml Nuclease-free Water. The buffer can be stored at -20°C for up to 2 weeks.

**1.1× lysis buffer**

Mix 220 µl 1 M Tris-HCl (pH8.0), 44 µl 5 M NaCl, 440 µl 10% SDS and 19.3 ml Nuclease-free Water for 20 ml 1.1× lysis buffer. Sterilize through 0.22 µm filter, store at room temperature (RT, 20–25 °C) for up to 1 month.

**Omni-ATAC RSB**

Mix 500 µl 1 M Tris-HCl (pH 7.5), 100 µl 5 M NaCl, 150 µl 1 M $MgCl_2$, and 49.25 ml Nuclease-free Water for 50 ml omni-ATAC RSB. Sterilize through 0.22 µm filter, stored at 4°C for up to 2 months.

**RSB-DTN**

Mix 487.5 µl Omni-ATAC RSB, 5 µl 10% Tween-20, 5 µl 10% IGEPAL CA-630 and 2.5 µl 2% Digitonin. Make fresh.

**RSB-T**

Mix 10 ml Omni-ATAC RSB and 100 µl 10% Tween-20. Store at 4°C for up to 1 month.

**0.1% digitonin**

Mix 0.5 µl 2% digitonin stock with 9.5 µl Nuclease-free Water. **CRITICAL** Make fresh and avoid freeze thaw cycles. The Promega 2% digitonin stock is dissolved in DMSO. Thaw the digitonin stock at room temperature.

**4× THS TD buffer**

Mix 132 µl 1 M Tris-HCl (pH8.0), 52.8 µl 5 M Potassium acetate, 40 µl 1 M Magnesium acetate, 640 µl N, N-Dimethylformamide (DMF) and 135.2 µl Nuclease-free Water for 1 ml 4× THS TD buffer. Store at -20°C for up to 6 months.

**2× Stop Buffer**

Mix 500 µl 1x DPBS, 40 µl 0.5M EDTA (pH=8.0), 66 µl 30% BSA and 394 µl Nuclease-free Water for 1 ml 2× Stop Buffer. Make fresh.

**50% (w/w) PEG8000**

Mix 5 g PEG8000 and 5 ml Nuclease-free Water, sterilize through 0.22 µm filter, aliquot and store at 4°C for up to 2 months.

**5× NaCl RT Buffer**

Mix 1.25 ml Tris-HCl (pH =8.0), 375 µl 5 M NaCl, 75 µl $MgCl_2$, 250 µl 1 M DTT, 3.05 ml Nuclease-free Water to prepare 5 ml 5× NaCl RT Buffer. Sterilize through 0.22 µm filter, aliquot and store at -20°C for up to 2 months.

**80% (v/v) ethanol**

Mix 4 ml absolute ethanol with 1 ml Nuclease-free Water. Make fresh.

#### Procedures

**Prepare Tn5 transposome complex Timing 2.5 h**

1. Dissolve ME_S5, ME_S7 and ME_bottom oligos respectively in annealing buffer to a final concentration of 100 µM.
2. Mix 25 µl ME_S5 oligo (100 µM) with 25 µl ME_Bottom oligo (100 µM) in a PCR tube, and anneal in a thermocycler as follows: 98 °C for 3 min, and slowly cool down to 16 °C with a temperature ramp of −0.1 °C/s, to generate S5_adaptor (50 µM).
3. Similarly, mix 25 µl ME_S7 oligo (100 µM) with 25 µl ME_Bottom oligo (100 µM) in a PCR tube and anneal to form S7_adaptor (50 µM).
4. Dilute the annealed 50 µM S5_adaptor or S7_adaptor with Nuclease-free Water to 20 µM.
5. To assemble Tn5-S5/S7 transposome, add 12 µl S5_adaptor (20 µM), 12 µl S7_adaptor (20 µM), 48 µl purified Tn5 (0.5 µg/µl), and 88 µl coupling buffer to a 1.5 ml tube, mix thoroughly by gently pipetting up and down 10 times. Incubate at room temperature for 1 h, then store at -20 °C.
6. To assemble Tn5-S7/S7 transposome, add 24 µl S7_adaptor (20 µM), 48 µl purified Tn5 (0.5 µg/µl), and 88 µl coupling buffer to a 1.5 ml tube, mix thoroughly by gently pipetting up and down 10 times. Incubate at room temperature for 1 h, then store at -20 °C.

**PAUSE POINT** The assembled Tn5 transposome could be stored at -20°C for up to 6 months.

**Prepare ISSAAC-seq lysis plate Timing 30 min**

**CRITICAL** Be careful to avoid the solution splashing when handling plates, which may cause index contamination among different wells. Multiple plates can be prepared at the same time for convenience.

7. Take out the "scATAC 10 µM i7 index plate" (refer to our plate-based scATAC-seq workflow, Nat Protoc, PMID: 34282334) and thaw at room temperature.
8. Mix 9 volumes of 1.1× Lysis Buffer and 1 volume of Truseq_S5_short primer (100 µM in Nuclease-free Water) to reach a 10 µM concentration in 1× Lysis Buffer. Aliquot 1 µl to each well of a new 384-well plate. Label the plate as "ISSAAC lysis plate" and with the date.
9. Briefly centrifuge the thawed "scATAC-seq 10 µM i7 index plate" from above. Transfer 1 µl of 10 µM i7 index to the ISSAAC lysis plate using a 16-channel multichannel pipette.

10. Briefly centrifuge the ISSAAC lysis plate and seal the plate. Leave the plate on ice and carry on the experiment until the cells are ready. Alternatively, store the plate at -80 °C.

    CRITICAL Evaporation can still happen even in the -80 °C freezer due to occasional open and closing of the freezer door. Therefore, always check the liquid volume in the wells before carrying on experiments.

    PAUSE POINT The sealed plates can be stored at −80 °C for up to 1 year.

**Prepare nuclei from cell lines Timing 1 h**

CRITICAL Work quickly and always keep the samples on ice unless otherwise indicated.

CRITICAL All centrifugation should be performed at 4 °C in a swing bucket centrifuge. DO NOT use fixed-angle centrifuges, which will cause severe cell loss during washes.

CRITICAL When removing the supernatant, be careful not to disturb the cell pellet.

11. Coat the tube: add 0.5 ml 1× DPBS–0.5%BSA-RI (0.2 U/µl) to a 1.5 mL tube, invert the tube three to five times, then aspirate all liquid in the tube and inside the cap and discard.

    CRITICAL Cells may attach to the inner surface of the centrifugation tubes. Coating the tubes with 0.5% (w/v) BSA helps reduce cell loss when working with low amounts of cells.

12. Count cells using a C-Chip disposable hemocytometer and use Trypan Blue to check cell viability. Make sure more than 90% cells are viable.
13. Spin down 100,000 cells per sample, centrifuge at 500g, 4°C for 5 min.

    CRITICAL Perform all centrifugations on a swing bucket centrifuge to reduce cell loss.

14. Wash the pellet once with 500 µl ice-cold DPBS-0.5% BSA-RI, centrifuge at 500g, 4°C, 5 min.
15. Resuspend cells in 50 µl ice-cold RSB-DTN- RI (0.8 U/µl) and leave on ice for 3 min.
16. Add 1 ml ice-cold RSB-T to the tubes (1.05 ml in total), invert 3 times to mix.
17. Centrifuge at 1000g, 4 °C for 8 min to collect the nuclei.
18. Aspirate all supernatants and hold on ice.

**Chromatin tagmentation Timing 1 h**

19. Prepare 50 µl chromatin tagmentation reaction for each sample:

| | Stock | Final | Volume (50 µl) |
|---|---|---|---|
| 4×THS TD buffer | 4× | 1× | 12.5 µl |
| 0.1% Digitonin | 0.1% | 0.01% | 5 µl |
| Tn5-S5/S7 | - | - | 2.5 µl |
| RiboLock RNase inhibitor | 40 U/µl | 1.2 U/µl | 1.5 µl |
| SUPERaseIn RNase inhibitor | 20 U/µl | 0.4 U/µl | 1 µl |
| RnaseOUT RNase inhibitor | 40 U/µl | 0.8 U/µl | 1 µl |
| Nuclease-free Water | - | - | 26.5 µl |

CRITICAL Tn5 should be the last component to add, *i.e.,* add it right before use.

20. Resuspend each pellet in 50 µl tagmentation mixture by gently pipetting up and down 30 times.

    CRITICAL Perform gentle and slow pipetting to avoid disrupting the nucleus structure. Make sure that the nuclei are intact after pipetting. Mix gently and avoid bubbles.

21. Incubate at 30°C, 800 rpm for 30 min.
22. Add 50 µl 2× Stop buffer to each sample and gently pipette up and down 6~8 times to mix.
23. Centrifuge at 1000g, 4 °C for 5 min. Remove the supernatants and wash the pellet twice with 200 µl ice-cold 0.5×DPBS-0.5%BSA-RI.
24. Carefully remove all supernatants and hold on ice.

    CRITICAL The nuclei pellet may be invisible, be careful not to disrupt the nuclei at the bottom of the tube when aspirating the supernatant. It is okay to have 2–3 µl buffer leftover at this step.

**Reverse transcription Timing 1.5 h**

25. Prepare 100 µl RT reaction for each sample, slowly pipette up and down 10~20 times to mix.

| Reagent | Stock | Final | Volume (100 µl) |
|---|---|---|---|
| TruseqR1_oligo_dT | 10 µM | 2 µM | 20 µl |
| dNTP | 10 mM each | 0.5 mM each | 5 µl |
| Maxima H minus Reverse Transcriptase | 200 U/µl | 10 U/µl | 5 µl |
| RiboLock RNase Inhibitor | 40 U/µl | 2 U/µl | 5 µl |
| SUPERaseIn RNase Inhibitor | 20 U/µl | 0.2 U/µl | 1 µl |
| RnaseOUT RNase Inhibitor | 40 U/µl | 0.4 U/µl | 1 µl |
| 5× NaCl RT Buffer | 5× | 1× | 20 µl |
| 50% PEG8000 | 50% | 12% | 24 µl |
| Nuclease-free Water | - | - | 19 µl |

CRITICAL The 50% PEG8000 solution is viscous. Pipette slowly to ensure that no liquid remains along the tube sidewalls and tips.

26. Resuspend each pellet with 100 µl RT buffer by gently pipetting up and down 50 times.

    CRITICAL Perform gentle and slow pipetting to avoid disrupting the nucleus structure. Make sure that the nuclei are intact after pipetting. Mix gently and avoid bubbles.

27. Transfer the 100 µl RT mixture to a new PCR tube and incubated in a thermocycler for 10 min at 50˚C before cycling for three times at 8˚C for 12s, 15˚C for 45s, 20˚C for 45s, 30˚C for 30s, 42˚C for 2 min, and 50˚C for 3 min, followed by a final step at 50˚C for 5 min.

28. Transfer the RT mixture to a new, BSA coated and chilled 1.5 ml tube.

29. Centrifuge at 1000g, 4 °C for 5 min. Remove the supernatants and wash the pellet twice with 200 µl ice-cold 0.5×DPBS-0.5%BSA.

30. Carefully remove all supernatants and hold on ice.

    CRITICAL After reverse transcription, the nuclei will be partially lysed and the number of intact nuclei will decrease. The pellet may be invisible, be careful not to disrupt the nuclei at the bottom of the tube when aspirating the supernatant. It is okay to have 2–3 µl buffer leftover at this step.

**mRNA-cDNA hybrid tagmentation Timing 1 h**

31. Set up RNA-DNA hybrid tagmentation mix, prepare 50 µl for each sample:

| | Stock | Final | Volume (50 µl) |
|---|---|---|---|
| 4×THS TD buffer | 4× | 1× | 12.5 µl |
| 0.1% Digitonin | 0.1% | 0.01% | 5 µl |
| Tn5-S7/S7 | - | - | 1.5 µl |
| Nuclease-free Water | - | - | 31 µl |

32. Resuspend each pellet with 50 µl tagmentation buffer by gently pipetting up and down 30 times.

    **CRITICAL** Perform gentle and slow pipetting to avoid disrupting the nucleus structure. Make sure that the nuclei are intact after pipetting. Mix gently and avoid bubbles.

33. Incubate at 37°C, 800rpm, 30 min.
34. Add 50 µl 2× Stop Buffer to each sample, mix thoroughly by gentle pipetting 6~8 times.
35. Centrifuge at 1000g, 4 °C for 5 min. Remove the supernatants and wash the pellet twice with 200 µl ice-cold 0.5×DPBS-0.5%BSA.
36. Carefully remove all supernatants and hold on ice.

    **CRITICAL** The pellet may be invisible, be careful not to disrupt the nuclei at the bottom of the tube when aspirating the supernatant. It is okay to have 2–3 µl buffer leftover at this step.

**EXO I digestion and Gap fill-in Timing 30 min**

37. Prepare 50 µl reaction for each sample:

| Reagent | Stock | Final | Volume (50 µl) |
|---|---|---|---|
| dNTP | 10 mM each | 0.5 mM each | 2.5 µl |
| Maxima H Minus Reverse Transcriptase | 200 U/µl | 8 U/µl | 2 µl |
| Exo1 | 20 U/µl | 2 U/µl | 5 µl |
| 5× NaCl RT Buffer | 5× | 1× | 10 µl |
| Nuclease-free Water | - | - | 30.5 µl |

38. Resuspend each pellet in 50 µl reaction by gently pipetting up and down 30 times.

39. Incubate at 37°C for 15 min.
40. Centrifuge at 1000g, 4 °C for 5 min, aspirate all supernatants.
41. Add 600 μl ice-cold 0.5×DPBS-0.5%BSA to each sample, resuspend pellet and transfer to a FACS tube.
42. Stain with DAPI: Mix 1 μl DAPI stock solution (1 mg/ml; protect from light) with 9 μl $ddH_2O$, and then add 1.5 μl of diluted DAPI solution to the 600 μl sample, vortex briefly and hold on ice.

**FACS sorting Timing >30 min**

43. Take out the "ISSAAC lysis plate" and thaw at room temperature. Briefly centrifuge to collect the liquid back to the bottom of each well.
44. Load the DAPI-stained sample and sort one DAPI positive nucleus to every well of the ISSAAC lysis plate except the well P24 which is left as a no cell control.
45. After sorting, seal the plate immediately, and centrifuge at 1,000 g for 1 min.

    **CRITICAL** It is very important to centrifuge the plate to make sure the nuclei go down to the lysis buffer. Perform the centrifugation immediately after sorting every plate, which helps avoid losing the cells on the plate wall. (Refer to PMID: 34282334 for more suggestion on FACS optimization.)

    **PAUSE POINT** After centrifugation, the plate containing the sorted nuclei can be stored at −80 °C for up to 3 months.

**Pre-amplification and well barcodes addition Timing 1 h**

46. Nuclei lysis: Incubate the plate at 65°C for 15 min, then centrifuge at 1000g for 1 min.
47. SDS quenching: carefully remove the sealing film, add 1 μl 10% Tween-20 to each well, then centrifuge the plate at 1000g for 1 min.

    **CRITICAL** Be careful to avoid the solution splashing out of the wells.

    **CRITICAL** The presence of SDS in the lysis buffer may inhibit the following PCR. The addition of excess Tween-20 neutralizes the negative effect of SDS on PCR.
48. Add 3 μl NEB Q5 High-Fidelity 2× Master Mix to each well. The total volume of PCR reaction in each well by now is 6 μl. Tightly seal the plate, then centrifuge at 1000g for 1 min.
49. Perform RNA library pre-PCR for individual cells in 384-well plate:

| | Temperature | Time | Number of cycles |
|---|---|---|---|
| Gap fill-in | 72°C | 5 min | 1 |
| Initial denaturation | 98°C | 1 min | 1 |
| Denaturation | 98°C | 20 sec | 4 |
| Annealing | 63°C | 20 sec | |
| Extension | 72°C | 1 min | |
| Hold | 10°C | ∞ | 1 |

50. Centrifuge the plate at 1000g for 1 min.
51. Carefully remove the sealing film, add 1 µl NEB Q5 High-Fidelity 2× Master Mix and 1 µl Nextra_S5 short primer (10 µM in Nuclease-free Water) to each well. Tightly seal the plate, and then centrifuge at 1000 g for 1 min.
52. Perform ATAC+RNA library pre-PCR for individual cells in 384-well plate:

| | Temperature | Time | Number of cycles |
|---|---|---|---|
| Initial denaturation | 98°C | 1 min | 1 |
| Denaturation | 98°C | 20 sec | 6 |
| Annealing | 63°C | 20 sec | |
| Extension | 72°C | 1 min | |
| Hold | 10°C | ∞ | 1 |

53. Centrifuge the plate at 1000 g for 1 min, then carefully remove the sealing film.
54. Put a Low Dead Volume Reservoir on top of the 384-well plate, invert and centrifuge at 1000g for 1 min to pool all wells from a plate into the reservoir.
55. Transfer the pre-amplified PCR products to a new 15 ml tube.

    **CRITICAL STEP** Genomic fragments from each well is now barcoded by a unique i7 index. However, plate barcode has not been added yet at this stage. Different plates must be processed separately. DO NOT mix samples from different plates.
56. Purify the pre-amplified PCR products using Zymo DNA Clean &Concentrator kit and elute the products in 41 µl Nuclease-free Water. (Refer to PMID: 34282334 for demonstration of using an extender tube and a vacuum connector to purify large volume of reaction in a single column.)

**Exonuclease I digestion and purification Timing 1 h**

57. Prepare 50 μl EXO I digestion reaction for each sample to eliminate residual primers form pre-amplifications:

| Reagent | Stock | Final | Volume (50 μl) |
|---|---|---|---|
| 10× NEB 3.1 buffer | 10× | 1× | 5 μl |
| EXO I | | | 5 μl |
| Purified pre-products | - | - | 40 μl |

58. Incubate at 37°C for 30 min.
59. Inactivate EXO I by heating at 80 °C for 2 min.
60. Purify the product by 1.2× VAHTS DNA Clean Beads, and elute in 16 μl Nuclease-free Water.

#### Final library amplification and plate barcodes addition  Timing 1 h

61. Set up PCR reaction:

| | |
|---|---|
| 2.5 μl | ATAC_plate_S5xx (10 μM) |
| 5.0 μl | Illumina P7 (10 μM) |
| 2.5 μl | RNA_plate_S5xx (10 μM) |
| 25 μl | Q5 High-Fidelity 2× Master Mix |
| 15 μl | Purified DNA from above |

**CRITICAL** The ATAC_plate_S5xx and RNA_plate_S5xx primers have the ATAC/RNA sample barcodes. Use different primers for different samples if you want to sequence the samples in the same lane.

62. Perform PCR as follows:

<table>
<tr><th></th><th>Temperature</th><th>Time</th><th>Number of cycles</th></tr>
<tr><td>Initial denaturation</td><td>98°C</td><td>1 min</td><td>1</td></tr>
<tr><td>Denaturation</td><td>98°C</td><td>20 sec</td><td rowspan="3">8</td></tr>
<tr><td>Annealing</td><td>63°C</td><td>20 sec</td></tr>
<tr><td>Extension</td><td>72°C</td><td>1 min</td></tr>
<tr><td>Final extension</td><td>72°C</td><td>5 min</td><td></td></tr>
<tr><td>Hold</td><td>10°C</td><td>∞</td><td>1</td></tr>
</table>

63. Resulting library was purified with 1.0× VAHTS DNA Clean Beads and elute in 20 μl Nuclease-free Water.

**<u>Library QC and sequencing Timing 4~6 d</u>**

64. Measure the concentration of each library by Qubit according to the manufacturer's instructions.
65. Dilute the libraries to 1~10 ng/μl using Nuclease-free Water.
66. Check the library size distribution on an Agilent high-sensitivity chip. A broad peak with an average size of 200–1000 bp will be observed.
67. Sequence the libraries for 150 x 8 x 8 x 150 cycles on Novaseq 6000 platform (Illumina).

#### Timing

| | |
|---|---|
| Prepare Tn5 transposome complex | 2 h 30 min |
| Prepare ISSAAC-seq lysis plate | 30 min |
| Prepare nuclei from cell lines | 1 h |
| Chromatin tagmentation | 1 h |
| Reverse transcription | 1 h 30 min |
| mRNA-cDNA hybrid tagmentation | 1 h |
| EXO I digestion and Gap fill-in | 30 min |
| FACS sorting | > 30 min |
| Pre-amplification and well barcodes addition | 1 h |
| Exonuclease I digestion and purification | 1 h |
| Final library amplification and plate barcodes addition | 1 h |
| Library QC and sequencing | 4~6 d |

#### Anticipated Results

**<u>Step 64):</u>** The library concentration is usually around 5~30 ng/μl (depending on cell types), which is sufficient for sequencing.

**Step 66):** A broad peak with an average size of 200–1000 bp will be observed.

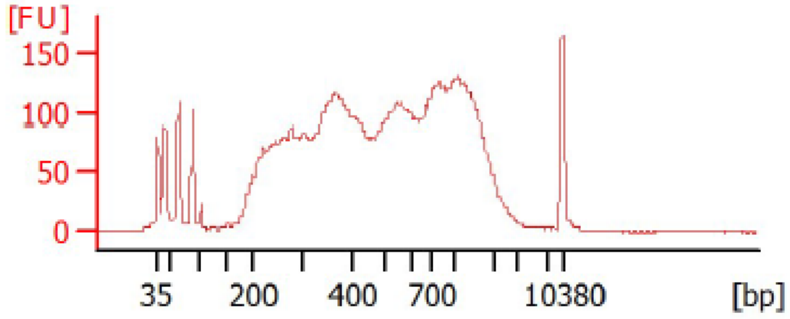
